# Collateral cleavage of 28s rRNA by RfxCas13d causes death of mice

**DOI:** 10.1101/2022.01.17.476700

**Authors:** Yunfei Li, Junjie Xu, Xuefei Guo, Zhiwei Li, Lili Cao, Shengde Liu, Ying Guo, Guodong Wang, Yujie Luo, Zeming Zhang, Xuemei Wei, Yingchi Zhao, Tongtong Liu, Xiao Wang, Huawei Xia, Ming Kuang, Qirui Guo, Junhong Li, Luoying Chen, Yibing Wang, Qi Li, Fengchao Wang, Qinghua Liu, Fuping You

**Author notes:** These authors contributed equally to this work (Yunfei Li, Junjie Xu, Xuefei Guo, Zhiwei Li). Correspondence (F.Y.) and (Q.L.).

## Abstract

The CRISPR-Cas13 system is an RNA-guided RNA-targeting system, and has been widely used in transcriptome engineering with potentially important clinical applications. However, it is still controversial whether Cas13 exhibits collateral activity in mammalian cells. Here, we found that knocking down gene expression using RfxCas13d in the adult brain neurons caused death of mice, which was not resulted from the loss of target gene function or off-target effects. Mechanistically, we showed that RfxCas13d exhibited collateral activity in mammalian cells, which is positively correlated with the abundance of target RNA. The collateral activity of RfxCas13d could cleave 28s rRNA into two fragments, leading to translation attenuation and activation of the ZAKα-JNK/p38-immediate early gene (IEG) pathway. These results provide new mechanistic insights into the collateral activity of RfxCas13d and warn that the biosafety of CRISPR-Cas13 system needs further evaluation before applying it to clinical treatments.

## Introduction

Clustered regularly interspaced short-palindromic repeats (CRISPR) and accompanying CRISPR-associated (Cas) proteins constitute the adaptive CRISPR-Cas immune system in bacteria and archaea, which protects the bacteria from invaders, including phages and mobile genetic elements. The defense process can be divided into three stages: Adaptation, incorporation of foreign DNA fragments into CRISPR array as spacers; CRISPR RNA (crRNA) biogenesis, CRISPR array is transcribed into a long precursor crRNA (pre-crRNA), and then processed into mature crRNAs; Interference, Cas effector proteins, under the guidance of crRNAs, specifically recognize and cleave foreign genetic elements. The rapid evolutionary arms race between bacteria and mobile genetic elements has greatly enriched CRISPR-Cas systems, which have been harnessed for various research and therapeutic applications. According to the structure and function of Cas effector proteins, CRISPR-Cas systems can be categorized into two classes, which are further subdivided into six types (type ◻-VI). Class 1 effectors comprise of multiple subunits, including type ◻, ◻, and ◻, while class 2 effectors are single large proteins, including type ◻, and ◻^[1]^. Due to their simplicity, Class 2 CRISPR-Cas systems have been widely developed as genome editing and transcriptional regulating tools, such as DNA-targeting Cas9 and Cas12, RNA-targeting Cas13.

Cas13 was originally found by mining microbial genome sequencing data using the highly conserved adaption protein Cas1 as the anchor^[2]^. Protein sequence alignments revealed that Cas13 contains two higher eukaryotes and prokaryotes nucleotide-binding (HEPN) domains and is predicted to possess ribonuclease (RNase) activity^[2]^. It was confirmed by subsequent experiments that Cas13-crRNA complex recognizes and cleaves the target RNA via base pairing between the crRNA and the target RNA^[3]^. Surprisingly, binding of the target RNA to Cas13-crRNA complex also activates a nonspecific RNase activity, which promiscuously cleaves bystander RNAs without complementarity to the crRNA, leading to cell death or dormancy in bacteria^[3, 4]^. This activity was referred as collateral activity of Cas13 and had been ingeniously developed as molecular diagnosis tool *in vitro*^[5]^. However, this collateral activity has not been detected in mammals. Theoretically, compared with Cas9-mediated gene knockout technology, Cas13 can accurately distinguish different transcripts of the same gene, and then study their function individually. Besides, Cas13-mediated gene silencing does not change genomic DNA, so this gene silencing is reversible and considered safer than Cas9, which has advantages over Cas9 in the treatment of some acquired diseases. Moreover, accumulating evidence over the past decade highlights that noncoding RNAs play important roles in various cellular processes. Cas13 is more suitable for noncoding RNAs research than Cas9.

Currently, there are six subtypes identified in the Cas13 family, including Cas13a, Cas13b, Cas13c, Cas13d, Cas13X and Cas13Y^[2, 6–10]^. Since their discovery, Cas13’s subtypes, such as LwaCas13a, PspCas13b, RfxCas13d and Cas13X.1, have been widely used in knockdown experiments in mammalian cells, exhibiting higher efficiency and specificity than traditional RNA interference, and no collateral activity was detected^[3, 6, 8, 10]^. Among these Cas13’s subtypes, due to its advantages in efficiency and size, RfxCas13d was applied in mammals via adeno-associated virus (AAV) delivery, with no side-effects reported^[11–13]^. But several research groups have different opinions about collateral activity of Cas13. Kang group first reported that the collateral activity of LwaCas13a occurred in U87 cells, non-specifically cleaving non-target RNAs, leading to cell death^[14]^. Later, they and collaborators reported that this phenomenon also existed in HepG2, AT2, B16F10 and GL261 cells^[15, 16]^. Following their work, Gootenberg and Abudayyeh group reported that collateral activity was detected in U87 cells for LwaCas13a, PspCas13b and RfxCas13d, and in HepG2 and mES cells for RfxCas13d^[17]^. Yang group claimed that LwaCas13a, RfxCas13d and Cas13X.1 exhibited collateral activity when targeting transiently overexpressing mCherry, but not endogenous genes in 293T cells^[10]^. Whereas, these studies did not figure out what effect collateral activity of Cas13 has on mammalian cells. Therefore, it is still controversial whether collateral activity of Cas13 exists in mammalian cells. More importantly, the safety of applying Cas13 to treatment needs to be carefully evaluated in animal models.

Here, we found that mice died when using RfxCas13d to knock down genes in brain neurons. The death would occur when target genes were present and obviously knocked down, but was not due to loss of gene function or off-target effects, which narrows down to collateral activity of Cas13. Then, we proved that RfxCas13d exhibited collateral activity in mammalian cells, which is positively correlated with the abundance of target RNA. The collateral activity of RfxCas13d cleaved 28s rRNA into two fragments, leading to translation attenuation and activation of the ZAK α-JNK/p38-IEG pathway.

## Results

### Mice died when knocking down *Sik3-S* in neurons using RfxCas13d

A recent study identified a *Sleepy (Sik3^Slp/+^)* mouse strain, which carries a mutation in the gene encoding salt-inducible kinase 3 (SIK3), a member of the AMP-activated protein kinase (AMPK) family^[18]^. The *Sleepy (Sik3^Slp/+^)* mice exhibit over 4 h increase in daily non-rapid eye movement sleep (NREMS) time and constitutively elevated NREMS delta power relative to wild-type (WT) littermates^[18]^. Interestingly, the *Sik3* gene encodes multiple transcripts due to alternative splicing. Our recent study identified a new *Sik3-S* transcript encoding ~72 kDa short isoform of SIK3^[19]^(Fig. S1a). To investigate the role of SIK3-S in sleep regulation, RfxCas13d was leveraged to specifically knock down *Sik3-S* by taking advantage of its ability to distinguish different transcripts of same gene (Fig. 1a). We designed eight crRNAs targeting *Sik3-S* and examined their knockdown efficiency in N2a cells through RT-qPCR (Fig. S1b). Our results showed that, in collaboration with RfxCas13d, all eight crRNAs, especially crRNA 1 and 8, caused significant knockdown of *Sik3-S* expression (Fig. 1b). Transcriptome analysis revealed that *Sik3* was specifically down-regulated while almost all the other genes remained unchanged when RfxCas13d was co-transfected with *Sik3-S* crRNA 1 or 8, compared with non-targeting (NT) crRNA (Fig. 1c).

**Figure 1.**
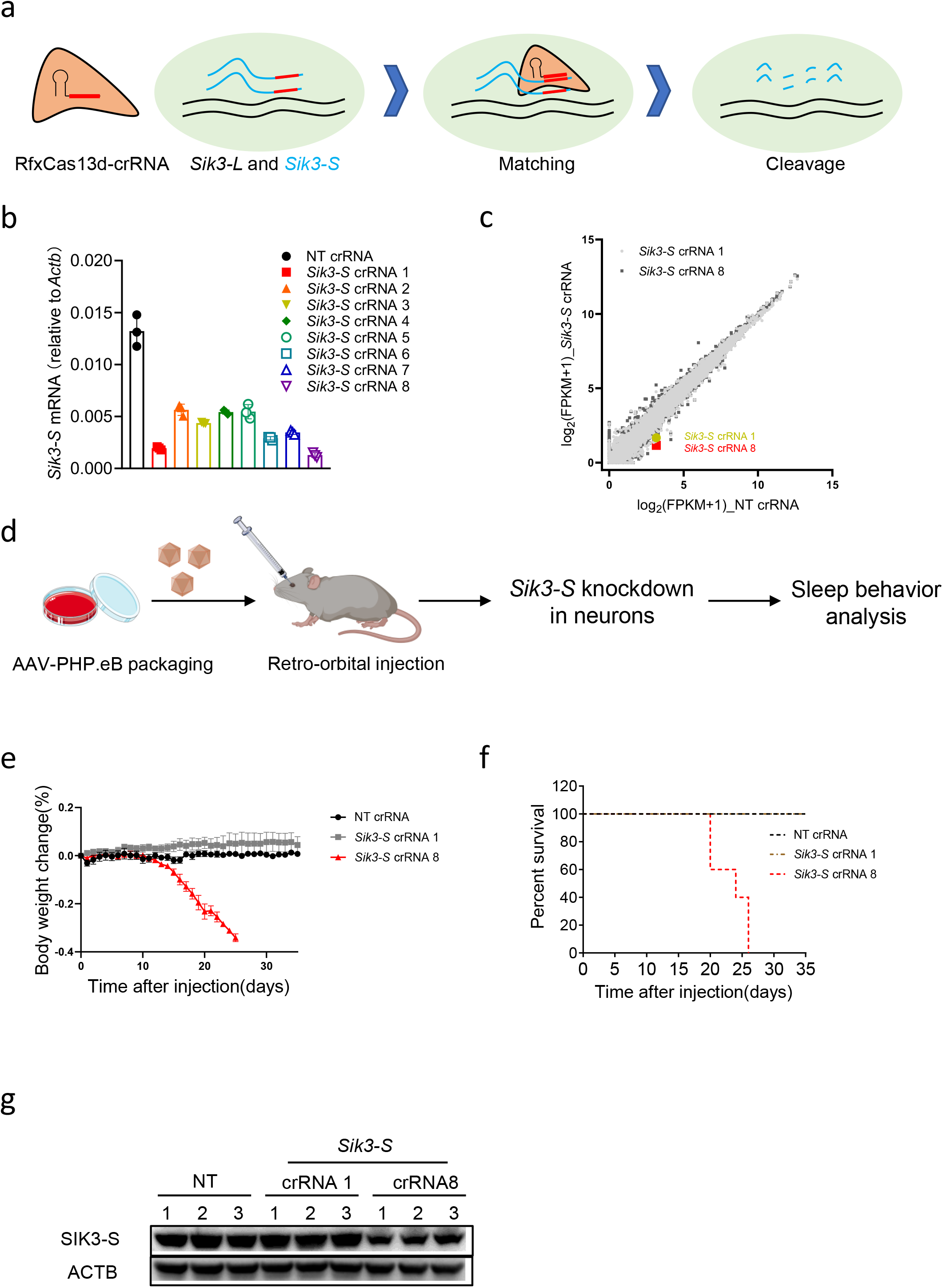
Mice died when knocking down *Sik3-S* in neurons using RfxCas13d. a. Schematic illustration of RfxCas13d-mediated specifically knockdown of *Sik3-S* not *Sik3-L* mRNA. *Sik3-L*: a transcript encoding a long isoform of SIK3. b. RT-qPCR to measure the knockdown efficiency of *Sik3-S* crRNAs. c. Transcriptome analysis in N2a cells 48 h post transfection of plasmids encoding RfxCas13d and *Sik3-S* crRNAs or NT crRNA. (n=3) d. Workflow of knocking down *Sik3-S* in the adult mouse brain neurons. e. Body weight change curve of ^LSL^RfxCas13d mice after injection of AAV-PHP.eB carrying *Sik3-S* crRNAs or NT crRNA. (n=5) f. Survival curve of ^LSL^RfxCas13d mice after injection of AAV-PHP.eB carrying *Sik3-S* crRNAs or NT crRNA. (n=5) g. Western blot to measure SIK-S expression level in brain lysates of ^LSL^RfxCas13d mice injected with AAV-PHP.eB carrying *Sik3-S* crRNAs or NT crRNA at 20 dpi.

We generated the ^LSL^RfxCas13d knock-in mice by inserting the CAG-loxP-STOP-loxP-RfxCas13d cassette into the *Rosa26* locus by homologous recombination (Fig. S1c). To knock down *Sik3-S* expression in the adult brain neurons, we retro-orbitally injected 12-week-old ^LSL^RfxCas13d adult mice with AAV-PHP.eB to deliver systemic expression of U6-driven crRNA and hSYN-driven Cre (Fig. S1d). AAV-PHP.eB could efficiently cross the blood brain barrier and transduced the majority of neurons and astrocytes across the adult mouse brain^[20]^. The human synapsin 1 gene promoter (hSYN) restricted the expression of Cre recombinase in neurons. Subsequently, Cre recombinase mediated excision of a tripartite transcriptional stop cassette (STOP) flanked by loxP to release the expression of RfxCas13d. RfxCas13d, under the guidance of *Sik3-S* crRNAs, specifically recognized and cleaved *Sik3-S* transcripts (Fig. 1d). Unexpectedly, mice injected with AAV-PHP.eB containing *Sik3-S* crRNA 8 began to lose weight at ~20 days post injection (dpi), and died at ~24 dpi (Fig. 1e-f). Brain lysates from *Sik3-S* crRNA 8 group, but not NT crRNA or *Sik3-S* crRNA 1 group, showed decreased SIK3-S expression, demonstrating that *Sik3-S* expression was specifically knocked down in this group (Fig. 1g). This phenomenon prevented our experiments from continuing, but aroused our curiosity-why mice died when knocking down *Sik3-S* using RfxCas13d in the adult mouse brain neurons.

### RfxCas13d mediated lethality was not due to the loss of target gene function

Several previous studies have proved that RfxCas13d can be used to knock down endogenous genes *in vivo* with no reported side-effects in liver, brain and eyes^[11–13]^. In addition, although the homozygous *Sik3* knockout mice can be created, but exhibit impaired chondrocyte during development, neonatal lethality and reduced size, indicating that *Sik3* is essential for mouse health and survival^[21, 22]^. Therefore, we first guessed whether the death of mice was caused by down-regulation of *Sik3-S*. To test this, we knocked out *Sik3* in the same neurons of mice by conventional Cre-loxP system. We generated Sik3-E5^flox^ mice by inserting two loxP sites into both sides of exon 5 in *Sik3* gene locus, and then delivered AAV-PHP.eB carrying hSYN-driven Cre into Sik3-E5^flox^ mice to knock out *Sik3*, with WT mice as control (Fig. S2a). At 21 dpi, brain lysates from Sik3-E5^flox^ mice showed lower SIK3-S expression level compared to WT mice (Fig. 2a). But Sik3-E5^flox^ mice behaved as normal as WT mice without loss of body weight or death (Fig. 2b-c, which means that knocking out *Sik3* in neurons will not cause mouse death. Thus, the mouse lethality that occurred when using RfxCas13d to knock down *Sik3-S* expression had nothing to do with the loss of functional SIK3-S.

**Figure 2.**
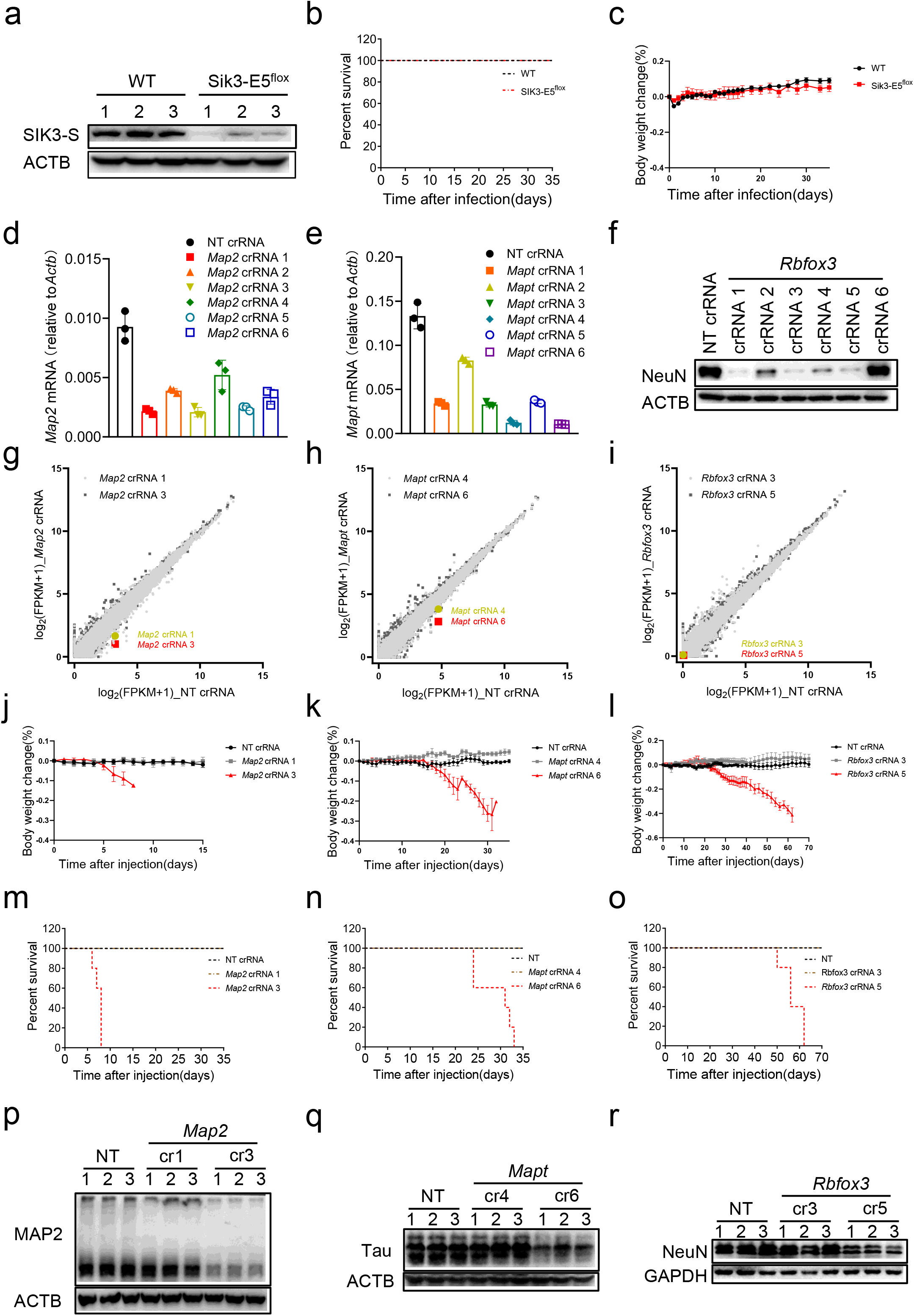
RfxCas13d mediated lethality was not due to the loss of target gene function. a. Western blot to measure SIK3-S expression level in brain lysates from Sik3-E5^flox^ and WT mice at 21 dpi. b. Survival curve of Sik3-E5^flox^ and WT mice after injection of AAV-PHP.eB-hSYN-Cre. (n=5) c. Body weight change curve of Sik3-E5^flox^ and WT mice after injection of AAV-PHP.eB-hSYN-Cre. (n=5) d-e. RT-qPCR to measure the knockdown efficiency of Mapt and Map2 crRNAs in N2a cells. f. Western blot to measure the knockdown efficiency of Rbfox3 crRNAs by knocking down overexpressing NeuN in HEK293T cells using RfxCas13d. g-i. Transcriptome analysis in N2a cells 48 h after transfection of plasmids encoding RfxCas13d and crRNAs. (n=3) j-l. Body weight change curve of ^LSL^RfxCas13d mice after infection of PHP.eB carrying *Map2* crRNAs, *Mapt* crRNAs or *Rbfox3* crRNAs. (n=5) m-o. Survival curve of ^LSL^RfxCas13d mice after injection of AAV-PHP.eB carrying *Map2* crRNAs, *Mapt* crRNAs or *Rbfox3* crRNAs. (n=5) p-r. Western blot to measure MAP2 (at 6 dpi), Tau (at 25 dpi) or NeuN (at 50 dpi) expression level in brain lysates of ^LSL^RfxCas13d mice after injection of AAV-PHP.eB carrying *Map2* crRNAs, *Mapt* crRNAs or *Rbfox3* crRNAs.

Map2, Tau and NeuN were well characterized neuron marker genes. Homozygous knockout of each of these genes did not lead to death of mice^[23–25]^. We thus selected them as targets and knocked down each of three targets in the same way as knocking down *Sik3-S in vivo*. In theory, when knocking down these genes individually using RfxCas13d in adult mouse brain neurons, the mice will not die due to the loss of these genes. We designed six crRNAs for each gene and tested their knockdown efficiency by RT-qPCR (Fig. 2d-e. Since N2a cells do not express NeuN, we tested the efficiency of crRNAs by knocking down expression of co-transfected NeuN plasmid in HEK293T (Fig. 2f). The two best crRNAs for each gene were selected for follow-up experiments. Transcriptome analysis showed that these three genes were specifically knocked down using corresponding crRNAs in tandem with RfxCas13d (Fig. 2g-i. Following the same protocol of knocking down *Sik3-S in vivo*, we knocked down these three genes respectively in ^LSL^RfxCas13d mice. Results showed that mice in *Map2* crRNA 3, *Mapt* crRNA 6 and *Rbfox3* crRNA 5 groups showed significant loss of body weight and death, while mice in the other groups behaved normally and survived (Fig. 2j-o. Besides, brain lysates showed that these three target genes were down-regulated in corresponding death groups (Fig. 2p-r). Taken together, these data suggested that RfxCas13d mediated mouse death was not due to the loss of target gene function.

### RfxCas13d mediated lethality was not caused by off-target effects

One ongoing concern using any CRISPR-Cas system for gene editing is off-target effects^[26]^. Thus, it’s necessary to determine whether RfxCas13d-mediated mouse death is caused by off-target effects. Firstly, Ai14 (Rosa-CAG-LSL-tdTomato-WPRE) reporter mice were introduced and crossed with ^LSL^RfxCas13d mice to generate ^LSL^RfxCas13d^+/fl^Ai14^+/fl^ mice^[27]^ (Fig. S3a-b. We designed seven crRNAs targeting tdTomato and tested their knockdown efficiency in N2a cells stabling expressing tdTomato by RT-qPCR. Among these crRNAs, crRNA 4 and 7 significantly damped the expression of tdTomato (Fig. 3a). Transcriptome analysis showed that tdTomato was specifically knocked down in cells transfected with tdTomato crRNA 4 or 7 (Fig. 3b). Moreover, the expression of the reported lethal genes was not affected by these two crRNAs. Then, we knocked down tdTomato in ^LSL^RfxCas13d^+/fl^Ai14^+/fl^ mice in the same way as knocking down *Sik3-S*. Results showed that mice injected with AAV-PHP.eB carrying tdTomato crRNA 7 began to lose weight at ~12 dpi and died at ~15 dpi, mice in the other groups behaved normally and survived (Fig. 3c-d). Brain lysates showed that tdTomato expression was lower in tdTomato crRNA 7 group than NT crRNA or tdTomato crRNA 4 group (Fig. 3e).

**Figure 3.**
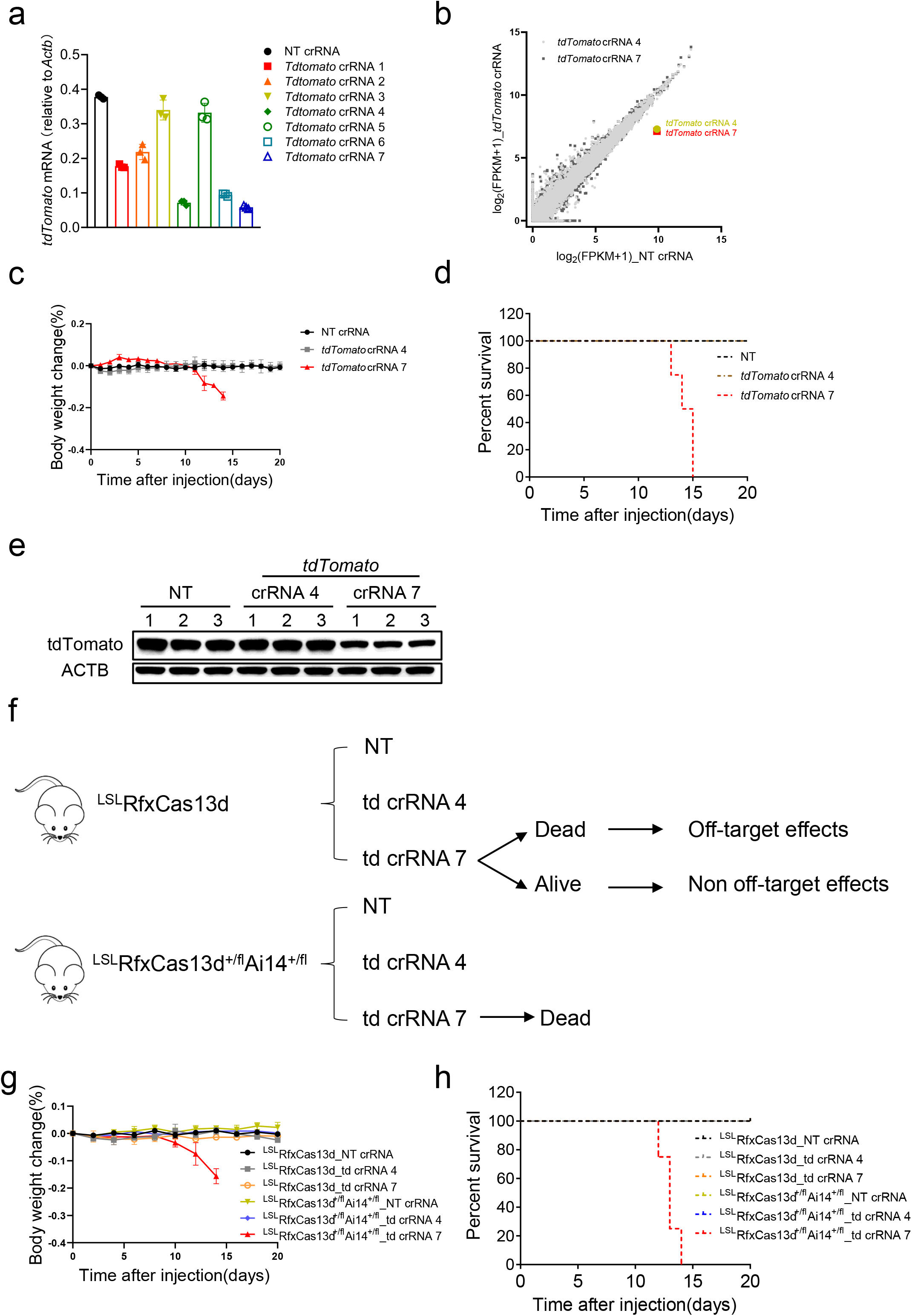
RfxCas13d mediated lethality was not caused by off-target effects. a. RT-qPCR to measure the knockdown efficiency of tdTomato crRNAs in N2a cells stably expressing tdTomato (N2a-td). b. Transcriptome analysis of N2a-td cells 48 h after transfection of plasmids encoding RfxCas13d and tdTomato crRNAs or NT crRNA. (n=3) c. Body weight change curve of ^LSL^RfxCas13d^+/fl^Ai14^+/fl^ mice after injection of AAV-PHP.eB carrying tdTomato crRNAs or NT crRNA. (n=4) d. Survival curve of ^LSL^RfxCas13d^+/fl^Ai14^+/fl^ mice after injection of AAV-PHP.eB carrying tdTomato crRNAs or NT crRNA. (n=4) e. Western blot to measure tdTomato expression level in brain lysates of ^LSL^RfxCas13d^+/fl^Ai14^+/fl^ mice injected with AAV-PHP.eB carrying tdTomato crRNAs or NT crRNA at 12 dpi. f. Schematic of delivering AAV-PHP.eB carrying tdTomato crRNAs or NT crRNA into ^LSL^RfxCas13d^+/fl^Ai14^+/fl^ and ^LSL^RfxCas13d mice, and the possible outcomes. g. Body weight change curve of ^LSL^RfxCas13d and ^LSL^RfxCas13d^+/fl^Ai14^+/fl^ mice after injection of AAV-PHP.eB carrying tdTomato crRNAs or NT crRNA. (n=4) h. Survival curve of ^LSL^RfxCas13d and ^LSL^RfxCas13d^+/fl^Ai14^+/fl^ mice after injection of AAV-PHP.eB carrying tdTomato crRNAs or NT crRNA. (n=4)

Next, we simultaneously injected AAV-PHP.eB carrying tdTomato crRNA 4, 7 or NT crRNA into ^LSL^RfxCas13d and ^LSL^RfxCas13d^+/fl^Ai14^+/fl^ mice. Theoretically, if RfxCas13d-mediated mouse death was caused by off-target effects, both ^LSL^RfxCas13d and ^LSL^RfxCas13d^+/fl^Ai14^+/fl^ mice injected with AAV-PHP.eB carrying tdTomato crRNA 7 would die. However, if RfxCas13d-mediated mouse death was not caused by off-target effects, only ^LSL^RfxCas13d^+/fl^Ai14^+/fl^ mice injected with AAV-PHP.eB carrying tdTomato crRNA 7 would die (Fig. 3f). Results showed that only ^LSL^RfxCas13d^+/fl^Ai14^+/fl^ mice injected with AAV-PHP.eB carrying tdTomato crRNA 7 began to lose body weight at ~12 dpi and died at ~15 dpi, mice in the other groups behaved normally (Fig. 3g-h). These data suggested that RfxCas13d-mediated mouse death was not caused by off-target effects. Moreover, since tdTomato is a foreign gene and has no function in brain neurons, this result further support that RfxCas13d mediated lethality had nothing to do with the loss of target gene function.

### The collateral activity of RfxCas13d was determined by the abundance of target RNA in mammalian cells

Above data ruled out the possibility that the death of mice was caused by the loss of target gene function or off-target effects. In addition, interestingly, only when the target genes were present and obviously knocked down, the mice would die. It was reminiscent of collateral activity of Cas13. Firstly, we verified findings in previous studies^[14, 17]^. We transfected *in vitro*-synthesized EGFP crRNA or NT crRNA into U87 cells stably expressing LwaCas13a and EGFP. The protein sequence of LwaCas13a and the sequence of crRNA we used are the same as previously reported^[14]^. Results showed that EGFP mRNA was significantly knocked down at 4 h and 8 h post transfection of EGFP crRNA, not NT crRNA (Fig. S4a). RNA denaturing gel electrophoresis of total RNA showed that 28s and 18s rRNAs were intact, and not cleaved into multiple bands as previous reported (Fig. S4b). Gootenberg and Abudayyeh group reported that LwaCas13a, PspCas13b and RfxCas13d exhibit collateral activity in U87 cells, thereby reducing cell viability^[17]^. But according to their results, transfection of plasmids encoding LwaCas13a, PspCas13b or RfxCas13d into U87 cells, regardless of with targeting or NT crRNA, would affect cell viability, indicating that cell viability changes have nothing to do with collateral activity. Therefore, it is still unclear whether collateral activity of Cas13 exists in mammalian cells.

We transiently transfected plasmids encoding RfxCas13d, tdTomato and crRNAs into HEK293T cells. Results showed that the protein and RNA level of RfxCas13d decreased, when it knocked down tdTomato under guidance of tdTomato crRNAs not NT crRNA (Fig. 4a-b. However, this phenomenon would not occur when there was no target gene expression or using catalytically dead RfxCas13d (dRfxCas13d) (Fig. 4a-b. This suggested that the collateral activity of RfxCas13d was activated to cleave its own mRNA when RfxCas13d-crRNA complex bound and cleaved tdTomato mRNA. Interestingly, changes in protein levels were more obvious than changes in RNA levels (this will be explained later). The same phenomenon was observed when knocking down *Sik3-S* (Fig. S4c-d. LwaCas13a and PspCas13b also exhibited similar characteristics (Fig. S4e-f. Yang group found that LwaCas13a, RfxCas13d and Cas13X.1 exhibited collateral activity when targeting transiently overexpressing mCherry, but not endogenous genes, using EGFP stably expressed in HEK293T as the indicator of collateral effects^[10]^, which gives us a hint that collateral activity of Cas13 may relates with the abundance of target RNA. Besides, in bacteria, Cas13-induced dormancy requires target RNA levels to exceed an expression threshold^[28]^. And *in vitro* experiments proved that collateral activity of Cas13 is positively correlated with the abundance of target RNA^[5, 29]^. To verify whether this correlation is also present in mammalian cells, a HEK293T cell line inducibly expressing tdTomato was constructed leveraging the tetracycline-controlled Tet-On inducible expression system, and then transfected with plasmids encoding RfxCas13d and crRNAs under different concentration doxycycline treatment (Fig. 4c). Results showed that RfxCas13d was negatively correlated with tdTomato at the expression level, under co-transfection of RfxCas13d and tdTomato crRNAs instead of NT crRNA (Fig. 4c). These data indicated that the collateral activity of RfxCas13d was triggered and positively correlated with the abundance of target RNA in mammalian cells, when targeting exogenous genes.

**Figure 4.**
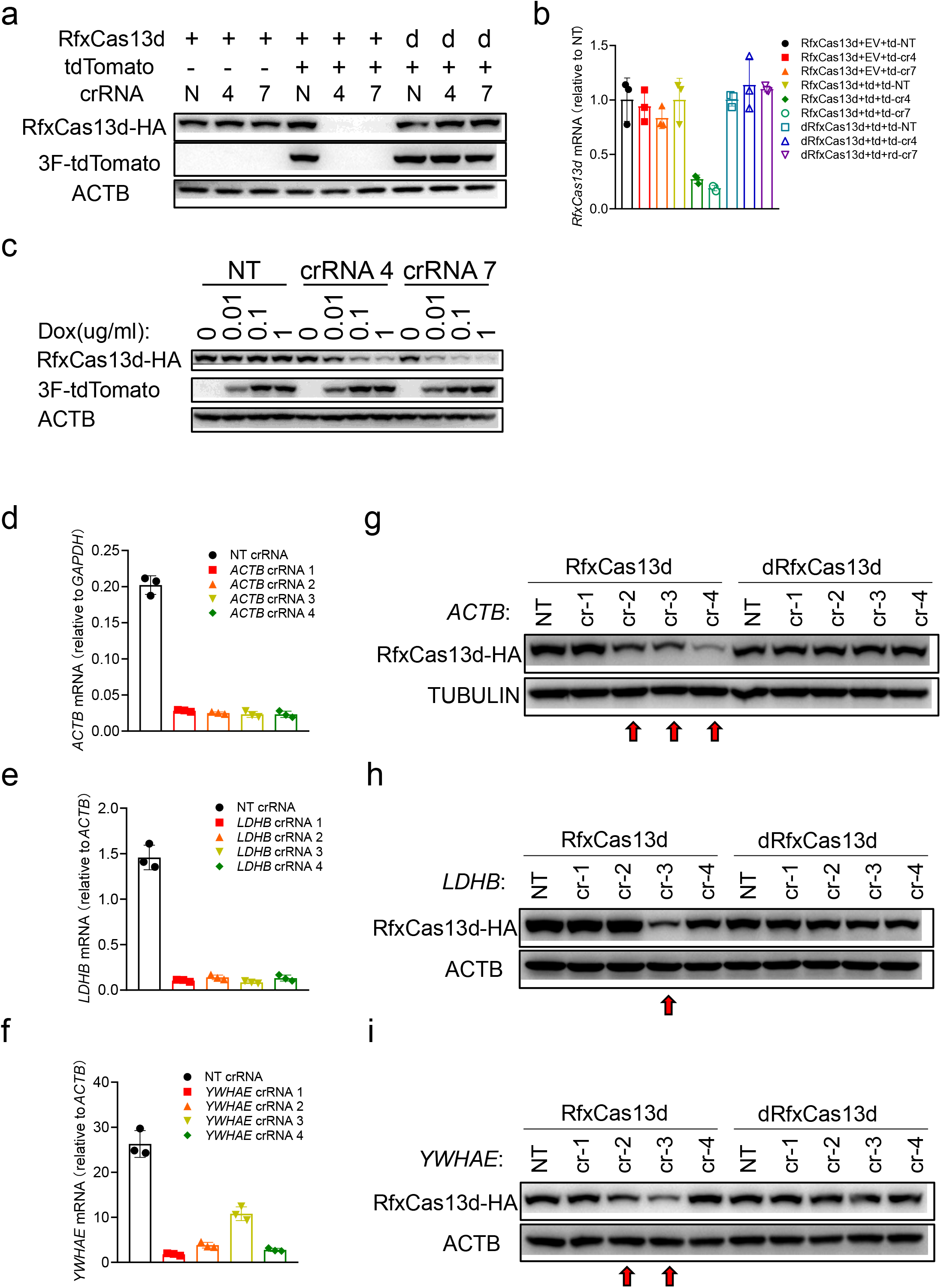
The collateral activity of RfxCas13d was determined by the abundance of target RNA in mammalian cells. a. Western blot to measure the expression level of RfxCas13d and tdTomato 24 h after transfection plasmids encoding RfxCas13d, tdTomato and crRNAs into HEK293T cells. 3F-tdTomato means tdTomato with 3x Flag tag at N-terminal. b. RT-qPCR to measure the RNA level of RfxCas13d in a. td represents tdTomato. EV represents empty vector. c. Western blot to measure the expression level of RfxCas13d and tdTomato 24 h after transfection of plasmids encoding RfxCas13d, tdTomato and crRNAs into inducible-expressing tdTomato HEK293T cells. Cells were treated with different doxycycline concentration in advance. d-f. RT-qPCR to measure the knockdown efficiency of ACTB, LDHB and YEHAE crRNAs in HEK293T cells. g-i. Western blot to measure the expression level of RfxCas13d/dRfxCas13d 24 h after transfection of plasmids encoding RfxCas13d/dRfxCas13d and crRNAs into HEK293T cells. cr-1/2/3/4: crRNA 1/2/3/4.

Next, we determined whether the collateral activity of RfxCas13d occurs when targeting endogenous genes in mammalian cells. We noticed that these endogenous genes previously used as targets are low in abundance^[8, 10]^. It is possible that collateral activity had been activated, but it was too weak to be detected. Therefore, we here selected several highly expressed genes as targets and designed four crRNAs for each gene. Then, plasmids encoding RfxCas13d/dRfxCas13d and crRNAs were transiently transfected into HEK293T cells. The knockdown efficiency of crRNAs was measured by RT-qPCR (Fig. 4d-f and S4g-j), and collateral activity was detected by measuring RfxCas13d expression level (Fig. 4g-i and S4k-n). Results showed that RfxCas13d, not dRfxCas13d, was down-regulated when targeting highly expressed genes, indicating that collateral activity was activated (crRNAs pointed by red arrows in Fig. 4g-i and S4k-n). Taken together, these data demonstrated that RfxCas13d exhibited collateral activity in mammalian cells, which is positively correlated with the abundance of target RNA.

### The collateral activity of RfxCas13d cleaves 28s rRNA into two fragments, leading to translation attenuation and activation of ZAKα-JNK/p38-IEG pathway

Although it was confirmed that the collateral activity of RfxCas13d existed in mammalian cells. It remains unknown whether and how this activity affects the biological process of cells. To this end, we constructed a HEK293T cell line stably expressing RfxCas13d (HEK293T-RfxCas13d) and then transfected with plasmids encoding target gene and corresponding crRNAs (Fig. S5a). In this way, RfxCas13d was fully expressed before the collateral activity was induced, which prevents the collateral activity of RfxCas13d from affecting its own expression by cleaving RfxCas13d mRNA (Fig. S5b). Thus, more RfxCas13d protein and the secondary induced collateral activity could be preserved than co-transfection. Then, cells were harvested 24 h post transfection for cell cycle distribution analysis, total RNA integrity analysis and RNA-seq (Fig. S5a). Analysis of total RNA integrity showed that, not only 28s and 18s rRNA, two additional bands but also were detected when co-transfecting of target genes and corresponding targeting crRNAs instead of NT crRNA into HEK293T-RfxCas13d cells (Fig. 5a). The same phenomenon occurred when targeting endogenous highly expression genes (Fig. S5c). In terms of size, these two additional bands looked like the products of 28s rRNA being cleaved. To test this, we did oligonucleotide extension assay to map cleavage sites (Fig. S5d). PCR and sanger sequencing revealed that 28s rRNA was cut into two fragments, one fragment of ~2100nt and the other segment of ~2800nt (Fig. S5e-f. Noticeably, several sequencing results detected until ~2187nt of 28s rRNA (marked by blue color), and one sequencing result revealed that a poly-A tail was added to 2187nt of 28s rRNA (marked by brown color) (Fig. S5f). There is “UU” behind 2187nt of the complete 28s rRNA (marked by red color) (Fig. S5f). And we proved that the collateral activity of RfxCas13d prefers to cleave poly-U *in vitro*, which is consistent with previous study^[30]^ (Fig. S5g-h. Therefore, this “UU” site (2188-2189nt) is likely to be the cleavage site by RfxCas13d on 28s rRNA (Fig. 5b). Those slightly shorter fragments may be caused by post-cleavage degradation. Interestingly, why did the collateral activity of RfxCas13d cleave 28s rRNA but not 18s rRNA? Theoretically, the abundance of 18s rRNA is also high, and there are also “UU” sites on it that can be cleaved. Besides, why did the collateral activity of RfxCas13d cut 28s rRNA at this “UU” site not others? We speculated that it may be due to the structure of RNA and RNA binding proteins (RBPs) that protect rRNAs from being cut. To test our speculation, we extracted total RNA from HEK293T cells and reconstituted the collateral activity of RfxCas13d *in vitro*, and founded that 28s and 18s rRNA were cleaved into multiple fragments (Fig. S5i), indicating that RNA structure and RBPs were involved in protecting RNA from the collateral activity of RfxCas13d. 28s rRNA is an important component of the ribosome. To determine whether the translation function of the ribosome was affected due to 28s rRNA breakage, SUnSET assay was employed to monitor protein synthesis. To avoid the impact of SIK3-S enzyme activity on translation, we used kinase dead SIK3-S (SIK3-S-K37M) instead of WT SIK3-S (Fig. S6a). Results showed that protein synthesis was attenuated when target genes were co-transfected with targeting crRNA, not NT crRNA (Fig. 5c). These data suggested that the collateral activity of RfxCas13d cleaved 28s rRNA into two fragments, thereby affecting the translation function of the ribosome. This may explain why in Fig. 4a, changes in protein levels of RfxCas13d were more obvious than changes in RNA levels (Fig. S5b). Cell cycle distribution analysis showed that co-transfection of target gene and targeting crRNAs, but not NT crRNA leaded to cell cycle arrest at G1 phase (Fig. 5d and S6b). This may be caused by impaired translation of protein.

**Figure 5.**
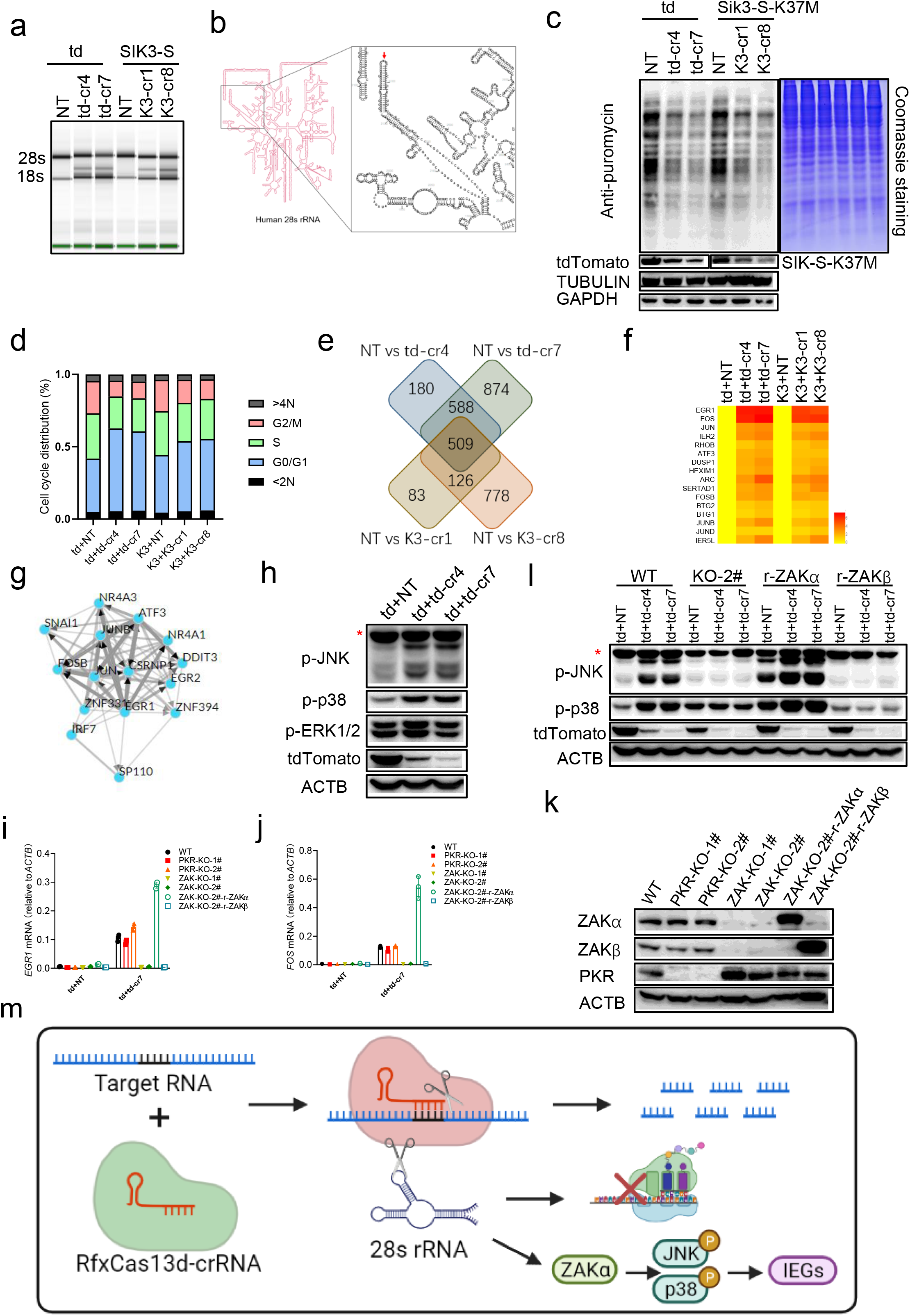
The collateral activity of RfxCas13d cleaves 28s rRNA into two fragments, leading to translation attenuation and activation of ZAKα-JNK/p38-IEG pathway. a. Total RNA of each sample was quantified by Agilent 2200 Bioanalyzer. td: tdTomato; td-cr4/7: tdTomato crRNA 4/7; K3-cr1/8: *Sik3-S* crRNA 1/8. b. Schematic illustration of the cleavage site of RfxCas13d on human 28s rRNA. c. SUnSET essay to measure the protein translation rate of HEK293T-RfxCas13d cells 24 h after transfection of plasmids encoding tdTomato/SIK3-S-K37M and corresponding crRNAs. d. Statistical diagram of cell cycle distribution in Fig. S6b. e. Schematic illustration of 509 common differentially expressed genes from four sets of comparisons. f. Heatmap to demonstrate the expression level of IEGs in different groups. log_2_(foldchange). g. Transcription factor enrichment analysis using ChEA3. There are Top 15 transcription factors. h. Western blot to measure the phosphorylation level of p38, JNK and ERK in HEK293T-RfxCas13d 24 h after transfection of plasmids encoding tdTomato and tdTomato crRNAs or NT crRNA. Red asterisk represents non-specific band. i-j. RT-qPCR to measure the RNA level of EGR1 and FOS in HEK293T-RfxCas13d cells 24 h after transfection of plasmids encoding tdTomato and tdTomato crRNAs or NT crRNA. k. Western blot to measure the expression level of ZAKα, ZAKβ and PKR in indicated cells. PKR-KO-1/2#: two strain of PKR knockout HEK293T-RfxCas13d cells; ZAK-KO-1/2#: two strain of ZAK knockout HEK293T-RfxCas13d cells; ZAK-KO-2#-r-ZAKα: re-expression of ZAKα in ZAK-KO-2#; ZAK-KO-2#-r-ZAKβ: re-expression of ZAKβ in ZAK-KO-2# l. Western blot to measure the phosphorylation level of p38, JNK 24 h after transfection of plasmids encoding RfxCas13d and tdTomato crRNA 7 or NT crRNA into indicated cells. Red asterisk represents non-specific band. KO-2#: ZAK-KO-2#; r-ZAKα: ZAK-KO-2#-r-ZAKα; r-ZAKβ: ZAK-KO-2#-r-ZAKβ. m. Schematic illustration that the collateral activity of RfxCas13d cleaves 28s rRNA into two fragments, leading to translation attenuation and activation of ZAKα-p38/JNK-IEGs pathway.

RNA-seq analysis showed that there were 509 common differentially expressed genes from four sets of comparisons (Fig. 5e). Interestingly, these were all up-regulated genes compared with NT crRNA (Supplementary Table 1). Among these genes, we noticed that multiple genes with obvious difference belong to IEGs (Fig. 5f). Besides, transcription factor enrichment analysis of 509 genes showed that multiple enriched transcription factors mediate the expression of IEGs, including JUNB, FOSB, JUN, EGR1, EGR2, ATF3, NR4A3, NR4A1 and CSRNP1 (Fig. 5g). IEGs are genes which are activated transiently and rapidly in response to various cellular stimuli. There are several pathways that lead to the activation of IEGs, such as the RhoA-actin and the ERK, JNK and p38 MAPK pathways^[31]^. We used inhibitors of these pathways to block IEGs expression, and found that IEGs expression can be blocked by p38 and JNK inhibitors not MEK1/2 or RhoA/C inhibitors (Fig. S6c-d. The combination of p38 and JNK inhibitors worked better (Fig. S6c-d. Consistently, western blot revealed increased phosphorylation of p38 and JNK, but not ERK1/2 (Fig. 5h), demonstrating that JNK and p38 were responsible for the expression of IEGs. Previous studies reported that ZAKα, the long isoform of ZAK, senses ribotoxic stress caused by rRNA damage or ribosome impairment, and then activates p38 and JNK pathways^[32]^. We speculated that ZAKα may sense 28s rRNA breakage caused by RfxCas13d and activate JNK and p38 pathways. To test this, we firstly used two ZAK inhibitors (6p^[33]^ and HY180) to block IEGs expression. IEGs expression can be inhibited by both inhibitors in a dose-dependent manner (Fig. S6e-f. Then, we knocked out ZAK in HEK293T-RfxCas13d cells and found that IEGs expression was blocked in ZAK knockout cells not PKR (another ribotoxic stress sensor) knockout cells (Fig. 5i-k. Besides, re-expression of ZAKα not ZAKβ (the short isoform of ZAK) in ZAK knockout cells can rescue the expression of IEGs (Fig. 5i-k. Consistently, western blot demonstrated that phosphorylation of p38 and JNK was blocked in ZAK knockout cell, and can be rescued by re-expression of ZAKα not ZAKβ (Fig. 5l). These data proved that ZAKα sensed 28s rRNA breakage caused by RfxCas13d and mediated phosphorylation of p38 and JNK, then activating IEGs expression. Taken together, these data demonstrated that the collateral activity of RfxCas13d cleaves 28s rRNA into two fragments, leading to translation attenuation and activation of the ZAKα-JNK/p38-IEG pathway (Fig. 5m).

## Discussion

Here, we initially utilized RfxCas13d to specifically knock down *Sik3-S* in the adult mouse brain neurons for studying its role in sleep. Unexpectedly, mice died when SIK3-S was obviously knocked down, which arises our curiosity about the death of mice. Subsequent *in vivo* experiments ruled out the possibility that RfxCas13d-mediated mouse death was due to loss of gene function or off-target effects, and demonstrated that mice would die when target genes were present and obviously knocked down. These data reminded us whether the death of mice was caused by activation of collateral activity when RfxCas13d recognized and cleaved target genes. To prove this, we confirmed that RfxCas13d exhibits collateral activity in mammalian cells, which is related to the abundance of target genes. Then, we founded that the collateral activity of RfxCas13d cleaves 28s rRNA into two fragments, leading to translation attenuation and activation of ZAKα-JNK/p38-IEG pathway. In conclusion, we found that RfxCas13d exhibits collateral effects in mammalian cells, causing death in mice.

Previous studies used RfxCas13d to knock down endogenous genes *in vivo*, such as in brain glia, liver and eyes, without side-effect reported. But we observed mouse death when using RfxCas13d to knock down endogenous genes in brain neurons. This discrepancy in experimental outcome may be because neurons are more important than other kinds of cells and not easy to regenerate, so neurons are more sensitive to collateral activity, and animal’s performance is more obvious. The death of mice warned us that the safety of RfxCas13d needs to carefully evaluation in animal models before applying it to treatment.

During the research, we found that exogenous genes were more likely to be cleaved by the collateral activity of RfxCas13d than endogenous genes, which may be due to the fact that endogenous genes hold more comprehensive RNA structure and closer combination with RBPs than exogenous genes. Therefore, changes in the expression of exogenous genes, such as EGFP or mCherry, are more suitable as indicators of the collateral activity of RfxCas13d, but the cleavage of exogenous genes does not represent that endogenous genes will also be cleaved.

The cleavage of 28s rRNA was easily observed, due to its high abundance and important role in cells. But it is still unclear whether the collateral activity of RfxCas13d cuts other RNAs. We observed 509 common differentially expressed genes from four sets of comparisons. IEGs are only part of them. It is unclear why other genes were up-regulated, which may be the consequence of the collateral activity of RfxCas13d cleaving RNAs other than 28s rRNA. So, there is an urgent need to establish a method to detect all the cleavage sites of Cas13’s collateral activity, which is crucial for Cas13’s optimization in the future. When RfxCas13d recognizes and cleaves the target RNA, its collateral activity is activated to cleaves RfxCas13d mRNA and 28s rRNA, which in turn negatively regulates its own expression and collateral activity. Therefore, it is recommended to express Cas13 in cells in advance, and then to induce its collateral activity and study cleavage sites. Studying the cleavage mechanism of Cas13’s collateral activity will not only direct Cas13’s optimization in transcriptome engineering via reducing or removing collateral activity, but also inspire us to develop new applications in mammalian cells taking advantage of collateral activity.

## Acknowledgement

This work was supported by the National Key Research and Development Program of China (2016YFA0500300; 2020YFA0707800), the National Natural Science Foundation of China (31570891; 31872736; 32022028; 81991505), Peking University Clinical + × (PKU2020LCXQ009), the Peking University Medicine Fund (PKU2020LCXQ009) and a grant from Zhuhai Science and Technology Innovation Bureau (ZH22036302200063PWC to Z. Yin). Thanks Yichen Deng for help in FCAS. Thanks Prof. Xiaoyun Lu for providing ZAK inhibitors 6p and HY180.

## Author Contributions

Y.L., J.X., F.Y. and Q.L. conceived the study and analyzed the data. Y.L., J.X. and Z.L. performed most of the experiments. X.G. was responsible for RNA-seq analysis. Y.W., S.L. and L.C. provided advise and technical help. Q.L. was responsible for AAV preparation. S.W. generated ^LSL^RfxCas13d mice. Y.G., G.W., Z.Z., X.W., Y.Z., T.L., X.W., H.X, M.K., Q.G., J.L. and L.C. assisted in the molecular experiments. Y.L., F.Y. and Q.L. wrote and revised the paper. J.X, S.L. and L.C. helped with the paper revision.

## STAR⍰Methods

### Cell culture

HEK293T, N2a and U87 cells were obtained from ATCC. Cells were cultured in DMEM medium supplemented with 10% FBS (Gibco) and 100 U/ml Penicillin-Streptomycin in a humidified incubator at 37 °C with 5% CO_2_. Additional 1% glutamine for U87 cells.

### Animals

All animals care and use adhered to the Guide for the Care and Use of Laboratory Animals of the Chinese Association for Laboratory Animal Science. All procedures of animal handling were approved by the Animal Care Committee of Peking University Health Science Center (permit number LA 2016240). ^LSL^RfxCas13d and Sik3-E5^flox^ mice on a C57BL/6J background were generated by the Transgenic Animal Center, NIBS, Beijing, China. Ai14 reporter mice were purchased from The Jackson Laboratory. Wild-type mice were purchased from Department of Laboratory Animal Science of Peking University Health Science Center, Beijing, China. Mice were kept and bred in pathogen-free conditions.

### Plasmids construction

Plasmids used in this study were prepared by standard molecular biology techniques and coding sequences entirely verified. All the mutants were constructed by standard molecular biology technique. Each mutant was confirmed by sequencing.

### Reagents and antibodies

Polyethylenimine (PEI) (764582, Sigma-Aldrich) and jetPRIME (114-15, Polyplus) were used for transfection. *In vitro*-synthesized crRNAs were purchased from GenScript. Quenched fluorescent reporter RNA was purchased from General Biology. Inhibitors used in this study including the following: p38 inhibitor SB203580 (HY-10256, MCE); JNK inhibitor SP600125 (HY-12041, MCE); MEK1/2 inhibitor U0126 (HY-12031, MCE); RhoA/C inhibitor (S7719, Selleck). ZAK inhibitors 6P and HY180 are gift from Prof. Xiaoyun Lu, Jinan University. Antibodies used in this study include the following: anti-HA (Rabbit, H6908, Sigma-Aldrich); anti-HA (Mouse, self-made); anti-SIK3 (Rabbit, self-made); anti-β-Tubulin (Mouse, HC101, TransGen Biotech); anti-ACTB (Mouse, 60008-1-Ig, Proteintech); anti-GAPDH (Mouse, 60004-1-Ig, Proteintech); anti-NeuN (Rabbit, 26975-1-AP, Proteintech); anti-Tau (Rabbit, 10274-1-AP, Proteintech); anti-MAP2 (Rabbit, 17490-1-AP, Proteintech); anti-ZAK (Rabbit, 28761-1-AP, Proteintech); anti-PKR (Rabbit, 18244-1-AP, Proteintech); anti-p-p38 (Rabbit, 4511, CST); anti-p-JNK (Rabbit, 4370, CST); anti-p-ERK1/2 (Rabbit, ET1609-42, HUABIO); HRP-conjugated Affinipure Goat Anti-Rabbit IgG(H+L) (SA00001-2, Proteintech); HRP-conjugated Recombinant Rabbit Anti-Mouse IgG Kappa Light Chain (SA00001-1, Proteintech).

### AAV-PHP.eB packaging, purification and injection

AAV-PHP.eB was packaged in AAVpro 293T cells (632273, Clontech). PHP.eB (Addgene#103005), pHelper (240071-54, Agilent) and transfer plasmids were co-transfected to AAVpro 293T cells by PEI MAX (24765, Polysciences). Cells were harvested by cell lifter (70-2180, Biologix) 72 h post-transfection. The cell pellets were suspended in 1× Gradient Buffer (10 mM Tris-HCl pH=7.6, 150 mM NaCl, 10 mM MgCl_2_). Five repeated cycles of liquid nitrogen freezing, 37°C water bath thawing and vortex were used to lyse cell. Then ≥50 U/ml of Benzonase nuclease (E1014, Milipore) were added to cell lysates and incubated at 37°C for 30 min. Centrifuge the cell lysate at 21,130g for 30 min at 4°C and transfer the supernatant to a pre-build iodixanol (D1556, Optiprep) step gradients (15%, 25%, 40% and 58%) for ultracentrifugation purification. Vacuum centrifuge at 41,000rpm, 4°C for 4 h, the virus particles were in the layer of 40% iodixanol gradient. Purified virus were extracted from the 40% virus containing layer by needle and concentrated using Amicon filters (UFC801096, EMD) and formulated in sterile phosphate-buffered saline (PBS) supplemented with 0.01% Pluronic F68 (24040032, Gibco). Virus titers were determined by qPCR while a linearized AAV plasmid as a standard. 1x10^12^ vg/mouse AAV-PHP.eB were delivered into mice via retro-orbital injection.

### RNA extraction and reverse transcription quantitative real-time PCR (RT-qPCR)

Total RNA from cells were isolated using TRIzol reagent (DP424, TIANGEN). 1ug RNA was reverse transcribed using HiScript II Q RT SuperMix (R223-01, Vazyme). Levels of these indicated genes were analyzed by qPCR amplified using SYBR Green (Q311, Vazyme). Data shown are the relative abundance of the indicated mRNA normalized to ACTB or GAPDH. The primers were list in Table 1.

**Table 1.**
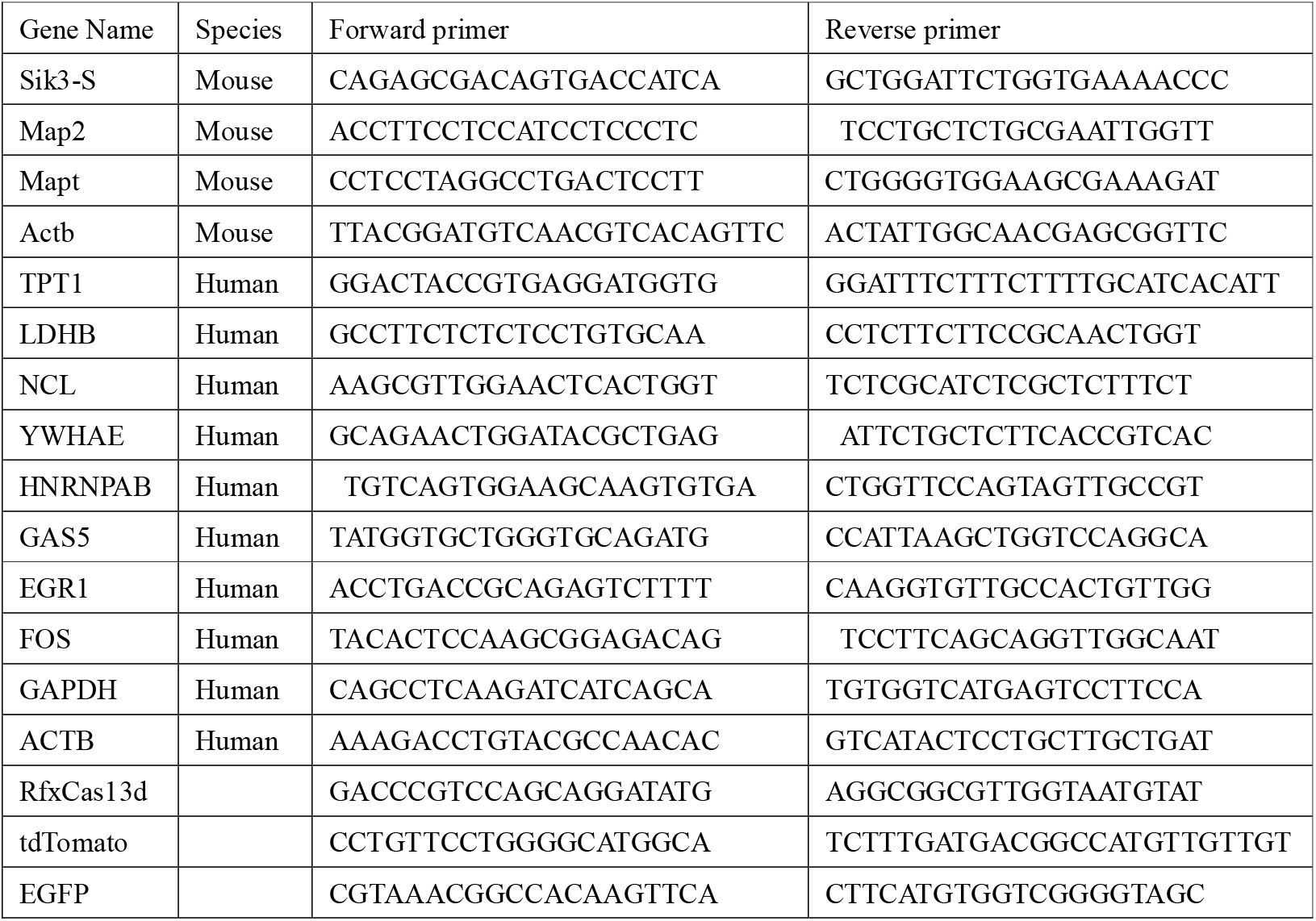
The sequence of qPCR primers used in this study.

### Measurement of crRNAs’ knockdown efficiency

Plasmids encoding RfxCas13d (addgene#109049) and crRNAs (addgene#109053) were transfected into N2a cells. 48 h after transfection, GFP-positive cells were sorted and collected through Fluorescence-Activated Cell Sorting (FACS), and then were extracted for total RNA. Then, levels of indicated genes were measured by RT-qPCR. All crRNAs used in this paper were listed in Table 2.

**Table 2.**
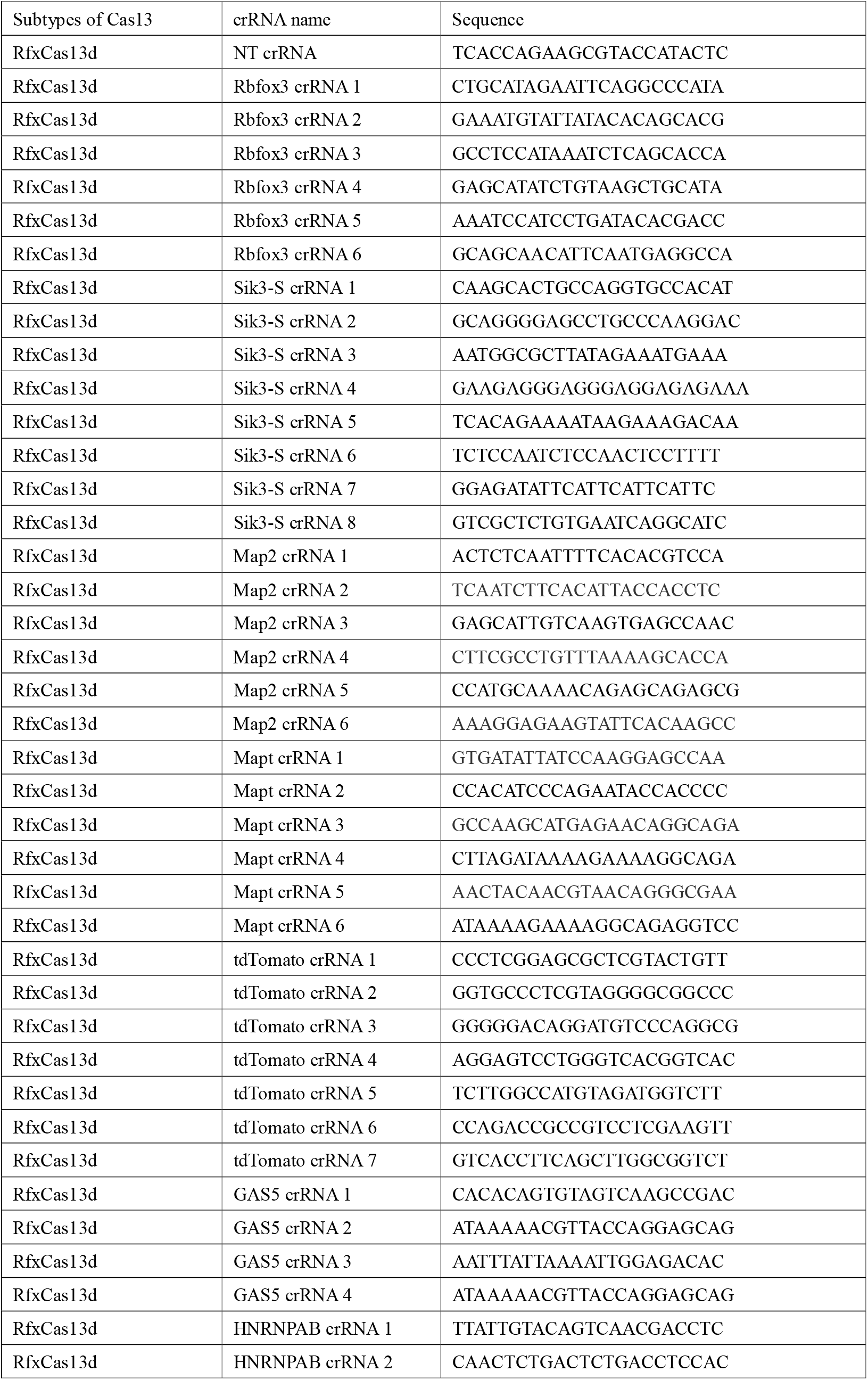
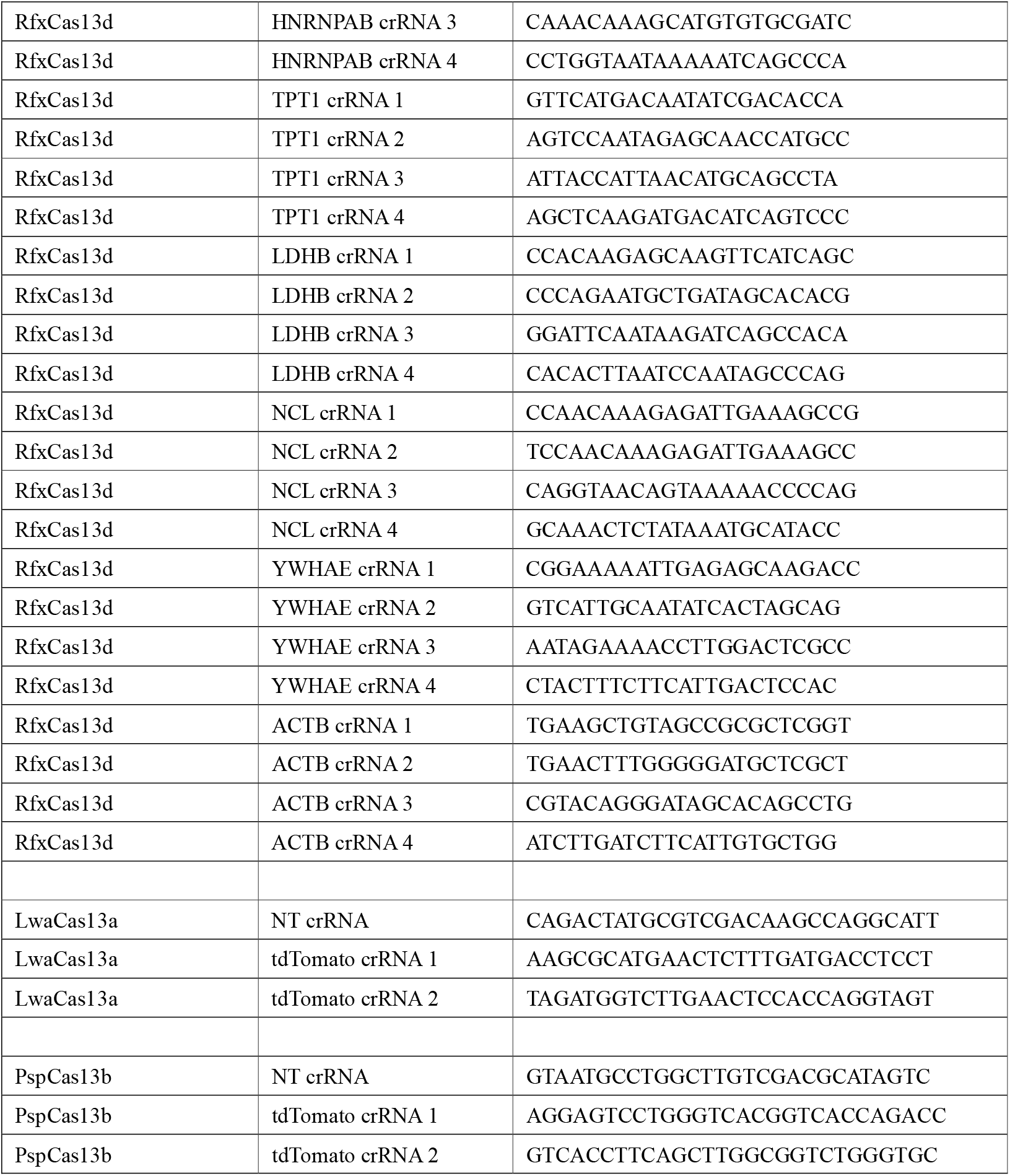
The sequence of crRNAs used in this study.

### Cell cycle distribution analysis

Cells were washed and collected using PBS to get rid of serum proteins at centrifugation at 1,200 rpm, 5 min. Resuspend pellet using precooling 70% EtOH solution to fix cells at least 30 min at 4°C. The cells can remain in this solution for up to one week. Dilute EtOH/cell suspension with PBS. Spin at 2,000-2,200 rpm for 10 min spin. Cells are much harder to pellet in EtOH. If EtOH is not diluted and the increased rate is not used, significant cell loss will be noticed. Wash cells three times using PBS and then stain cells using DAPI staining solution (C1005, Beyptime) for 30 min. Finally, cells were recorded by Fluorescence Activated Cell Sorting (FACS) and analyzed by FlowJo.

### Western Blotting

Cells were washed with PBS and lysed by incubation on ice for 10 min with RIPA lysis buffer (50 mM Tris, 150 mM NaCl, 0.1% SDS, 0.5% sodium deoxycholate, 1% Triton X-100, protease cocktail [C0001, Targetmol], and 1 mM PMSF). Brain tissue was firstly grinded in a mortar cooled on liquid nitrogen, and then lysed by incubation on ice for 30 min with RIPA lysis buffer. Supernatants were collected by centrifugation at 12,000 rpm for 10 min at 4°C, and then mixed up with loading buffer and boiled for 10 min. Samples were resolved by SDS/PAGE and transferred to 0.22 um nitrocellulose membrane (P-N66485, Pall). The membrane was blocked using skim milk for 30 min, then incubated overnight with primary antibodies, further incubated with the corresponding HRP-conjugated secondary antibodies and finally detected by enhanced chemiluminescence.

### SUnSET assay

Cells were incubated with puromycin (2.5 μg/ml) for 20 _ min and then washed with ice cold PBS and lysed using RIPA lysis buffer. Equal quantity of cell lysates was submitted to western blot using anti-puromycin antibody to detect protein synthesis. Signals were normalized with probing GAPDH and TUBULIN (loading control).

### Construction of Stable and inducible Expression Mammalian Cell Lines

For preparation of lentiviruses, HEK293T cells in 6-well plates were transfected with the lentiviral vector of interest (1,800 ng), the lentiviral packaging plasmids psPAX2 (600 ng) and pMD2.G (600 ng) and 12 ul of PEI (1 mg/ml). About 48 h after transfection, culture medium containing lentiviruses was collected and centrifugalized at 12,000 rpm for 10 min, and then filtered using 0.22 um filter. HEK293T, N2a and U87 cells were then infected at ~50% confluency by lentiviruses for 48 h, followed by selection with puromycin or hygromycin for 7 days. Monoclonal cells were obtained by limiting dilution.

### Oligonucleotide extension assay

Total RNA was ligated with oligonucleotide adaptor 1 or 2 respectively using T4 RNA Ligase 1 (M0204S, NEB) following manual. Then RNA was purified by ethanol precipitation and then reverse transcribed using R1 or R2 (R312-02, Vazyme). cDNA was amplified by PCR using F1+R1 or F2+R2. PCR products were firstly purified, then linked into T vector (CT101-01, Transgen Biotech), and finally sequenced by sanger sequencing.

The sequence of oligonucleotide adapters and primers:

adaptor 1:5-PO_4_-CTGTAGGCACCATCAATGGACCT-NH_2_-3 (DNA);

adaptor 2:5-NH_2_-CAGAAGGCACCAACAAAGGACC-OH-3 (RNA); F1: 5-ACCTGGGTATAGGGGCGAAAGAC-3 (DNA)

R1: 5-AGGTCCATTGATGGTGCCTACAG-3 (DNA)

F2: 5-CAGAAGGCACCAACAAAGGACC-3 (DNA) R2: 5-CCCTTAGAGCCAATCCTTATCCC-3 (DNA)

### Reconstitution of the collateral activity of RfxCas13d *in vitro*

To detect the collateral activity of RfxCas13d *in vitro*, we performed *in vitro* cleavage assay with 100 ng purified RfxCas13d protein, 100 ng synthesized tdTomato RNA, 100 ng crRNA, 2 μl RNase inhibitor (New England Biolabs), and 200 ng quenched fluorescent RNA reporters (6 nt polyA/U/G/C), in 100 μl reaction buffer (40 mM Tris-HCl, 60 mM NaCl, 6 mM MgCl_2_, pH 7.6)^[5]^. Reactions were incubated at 37°C for 1 h and measured the fluorescence of RNA reporters with microplate reader. In Fig. S5I, quenched fluorescent RNA reporters were replaced with total RNA extracted from HEK293T cells. Reactions were incubated at 37°C for 1 h and then RNA was purified by ethanol precipitation and quantified by Agilent 2200 Bioanalyzer.

### RNA denaturing gel electrophoresis

Make gel: Weigh 0.5 g of agarose powder, add it to 36.5 ml of DEPC water, and heat to completely dissolve the agarose. After cooling slightly (60-70°C), add 5 ml of 10x MOPS Running Buffer (C516042-0001, Sangon Biotech), 8.5 ml of 37% formaldehyde. Then pour the gel in the glue tank, insert the comb, and place it horizontally for use after solidification. Add samples: Mix the following reagents in a clean small centrifuge tube: 2 ul 10x MOPS Running buffer, 3.5 ul formaldehyde, 10 ul formamide (deionized), 4.5 ul RNA sample. Mix well, keep it at 60°C for 10 min, and cool quickly on ice. Add 3 ul of 10x loading buffer (B548318-0001, Sangon Biotech) and 0.5 ul of ethidium bromide, then mix well and add an appropriate amount to the sample well of the gel. Electrophoresis: Turn on the electrophoresis instrument, and stabilize the electrophoresis at 7.5 V/cm.

### Total RNA integrity analysis

Total RNA was extracted from cells and then quantified by Agilent 2200 Bioanalyzer.

### RNA-seq analysis

The sequencing data generated by illunima Noveseq PE150 in fastq file format was filtered by FastQC(https://www.bioinformatics.babraham.ac.uk/projects/fastqc/) and Trim-Galore(https://www.bioinformatics.babraham.ac.uk/projects/trim_galore/) softwares for quality control. Then the mouse genome version mm10 and the human genome version hg38 were used as reference genome to align the clean data with Subread software (http://subread.sourceforge.net/). The gene count matrix was calculated by the featureCounts (http://subread.sourceforge.net/) program. Then the gene count data was normalized using the FPKM formula. The differentially expressed genes were analysed by R package DESeq2 (https://bioconductor.org/packages/release/bioc/vignettes/DESeq2/inst/doc/DESeq2.html). Transcription factor enrichment analysis was conducted by ChEA3 (https://maayanlab.cloud/chea3/#top). The raw data and processed data were uploaded to the GEO Datasets (GSE193668).

### Statistical Analysis

The descriptive statistical analysis was performed with Prism version 8 (GraphPad Software). All data are presented as mean ± SEM. A two-tailed Student's t test assuming equal variants was used to compare two groups. In all figures, the statistical significance between the indicated samples and control is designated as *P < 0.05, **P < 0.01, ***P < 0.001, or NS (P > 0.05)

---

**Figure S1.**
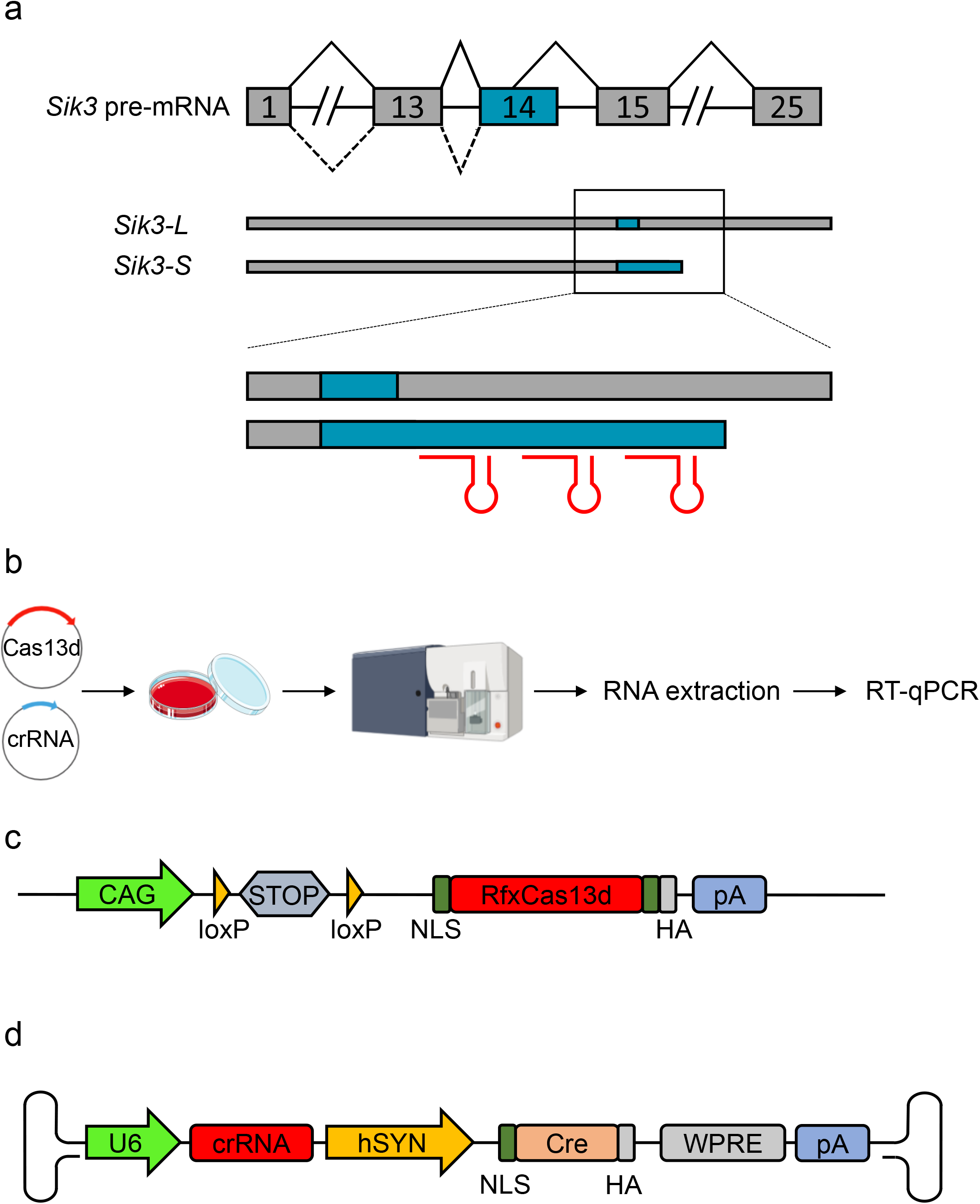
Mice died when knocking down *Sik3-S* in neurons using RfxCas13d. Linked to Figure 1. a. Schematic illustration of alternative splicing process to generate *Sik3-L* and *Sik3-S* transcripts, and design of *Sik3-S* crRNAs. b. Workflow of measuring the knockdown efficiency of crRNAs in N2a or HEK293T cells. c. Schematic diagram of ^LSL^RfxCas13d mice. NLS: nuclear localization sequence; HA: hemagglutinin tag; CAG: CAG promoter; U6: U6 promoter; STOP: a tripartite transcriptional stop cassette. d. Schematic illustration of AAV-PHP.eB plasmid used. hSYN: human synapsin 1 gene promoter.

**Figure S2.**
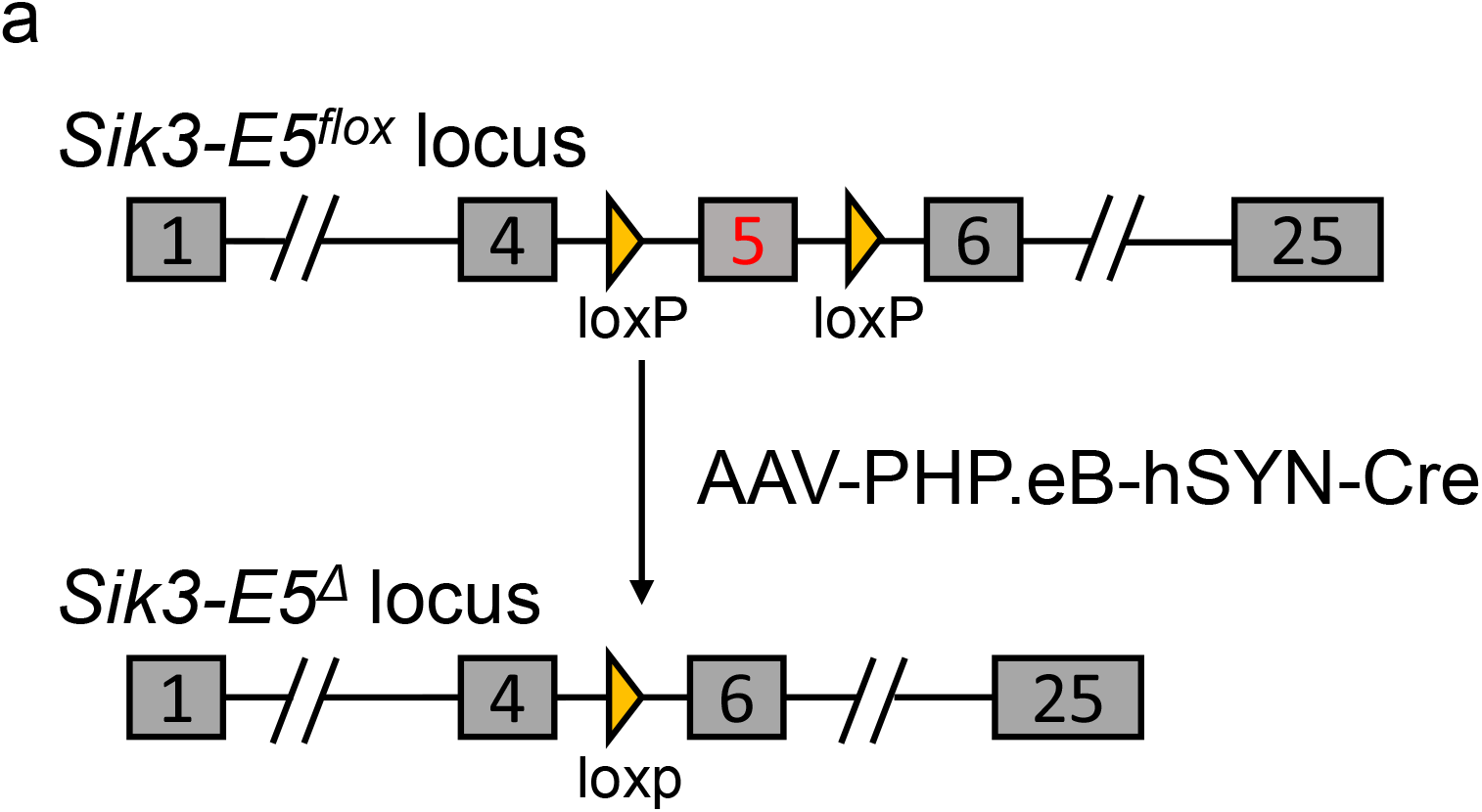
RfxCas13d mediated lethality was not due to the loss of target gene function. Linked to Figure 2. a. Schematic illustration of knocking out Sik3 by AAV-PHP.eB-hSYN-Cre injection of Sik3-E5^flox^ mice.

**Figure S3.**
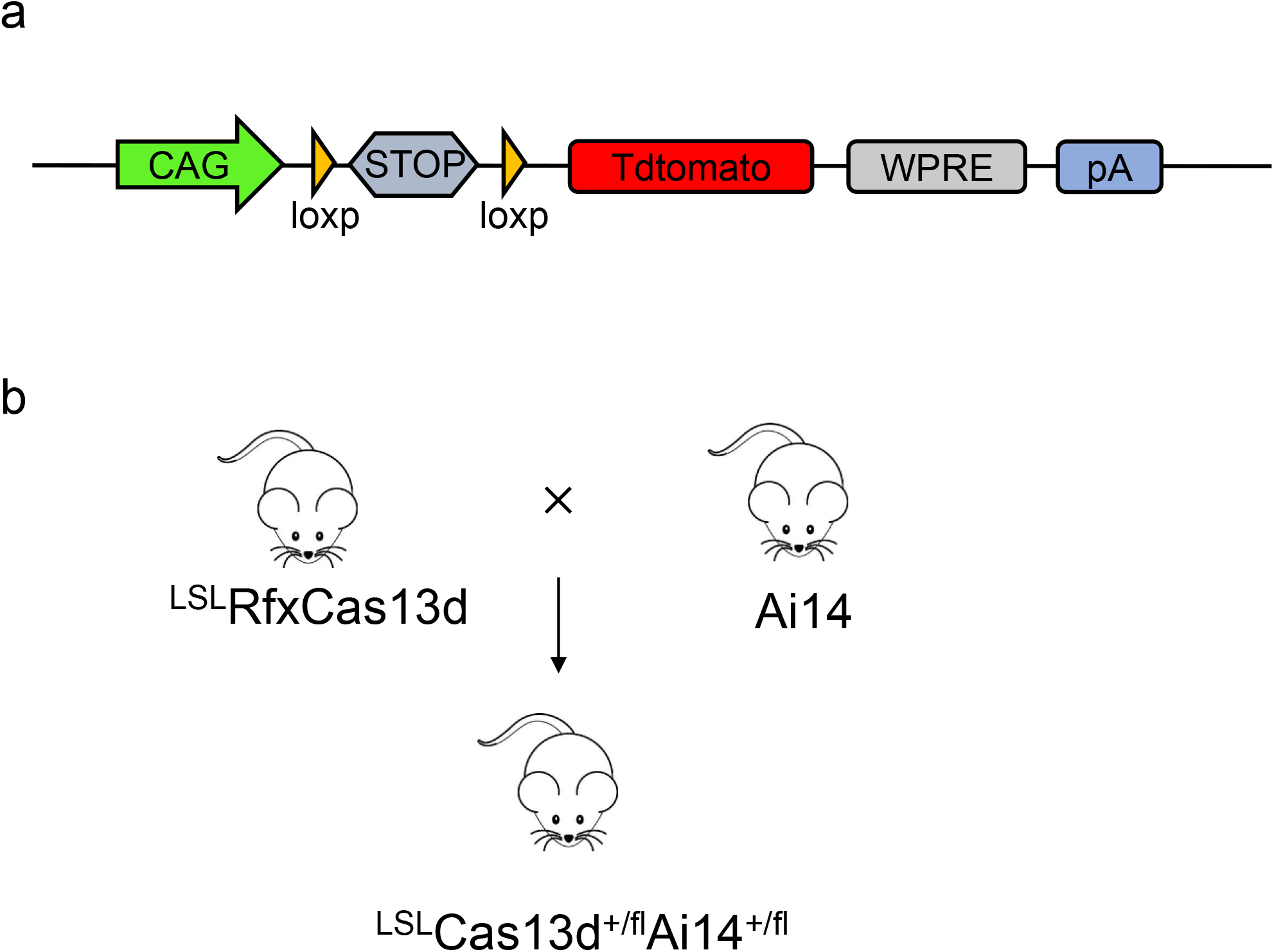
RfxCas13d mediated lethality was not caused by off-target effects. Linked to Figure 3. a. Schematic diagram of Ai14 reporter mice b. Schematic illustration of generating ^LSL^Cas13d^+/fl^Ai14^+/fl^ mice by crossing ^LSL^RfxCas13d mice with Ai14 mice.

**Figure S4.**
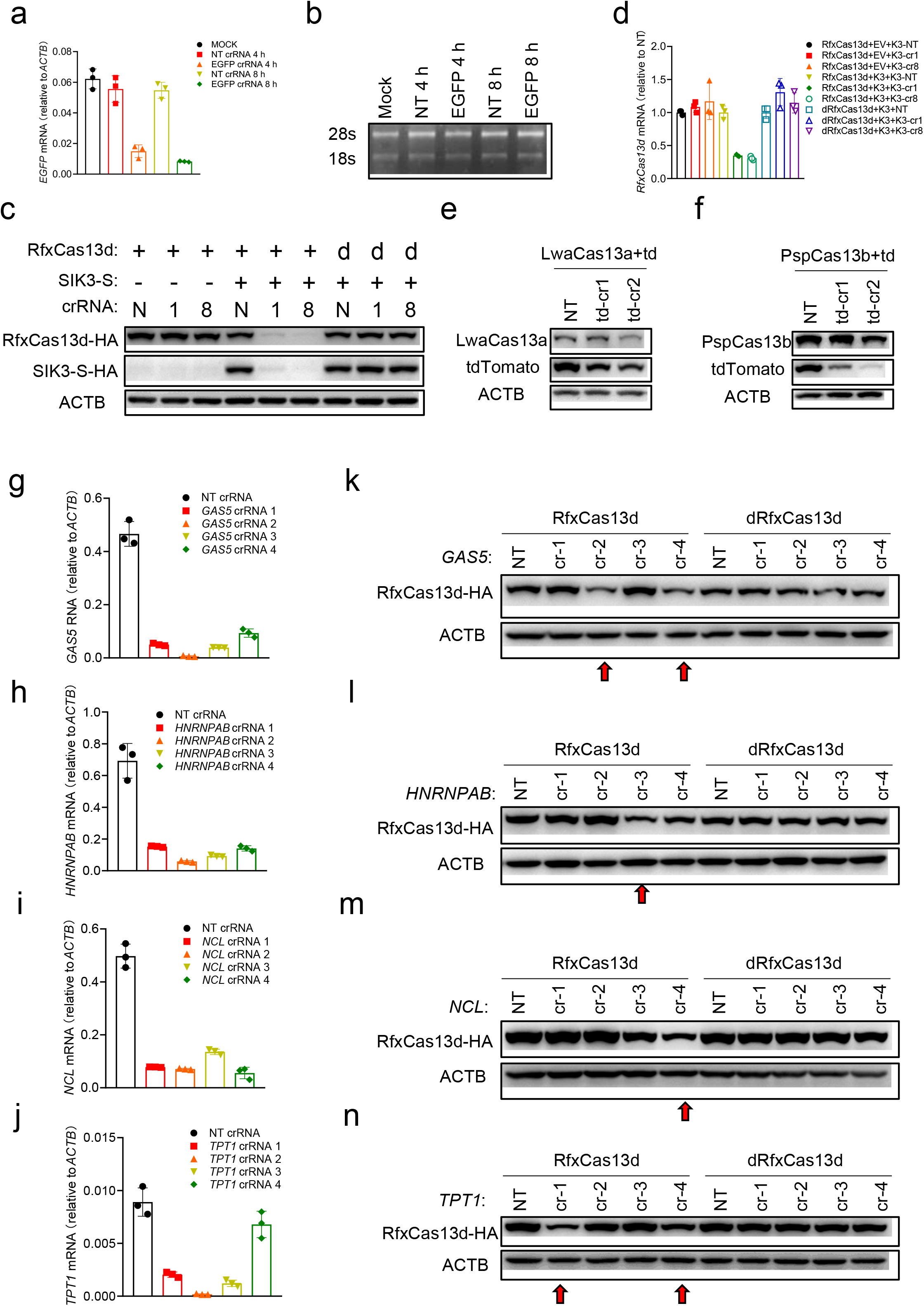
The collateral activity of RfxCas13d was determined by the abundance of target RNA in mammalian cells. Linked to Figure 4. a. RT-qPCR to measure the RNA level of EGFP after transfecting *in vitro*-synthesized EGFP crRNAs into U87 cells stably expressing LwaCas13a and EGFP. b. RNA denaturing gel to measure the integrity of total RNA in a. c. Western blot to measure the expression level of RfxCas13d and SIK3-S 24 h after transfection plasmids encoding RfxCas13d, SIK3-S and crRNAs into HEK293T cells. d. RT-qPCR to measure the RNA level of RfxCas13d in c. K3 represents SIK3-S; EV represents empty vector. e-f. Western blot to measure the expression level of LwaCas13a/PspCas13b and tdTomato 24 h after transfection of plasmids encoding LwaCas13a/PspCas13b, tdTomato and corresponding crRNAs into HEK293T cells. td-cr1: tdTomato crRNA 1; td-cr2: tdTomato crRNA 2. g-j. RT-qPCR to measure the knockdown efficiency of GAS5, HNRNPAB, NCL and TPT1 crRNAs in HEK293T cells k-n. Western blot to measure the expression level of RfxCas13d/dRfxCas13d 24 h after transfection of plasmids encoding RfxCas13d/dRfxCas13d and crRNAs into HEK293T cells. cr-1/2/3/4: crRNA 1/2/3/4.

**Figure S5.**
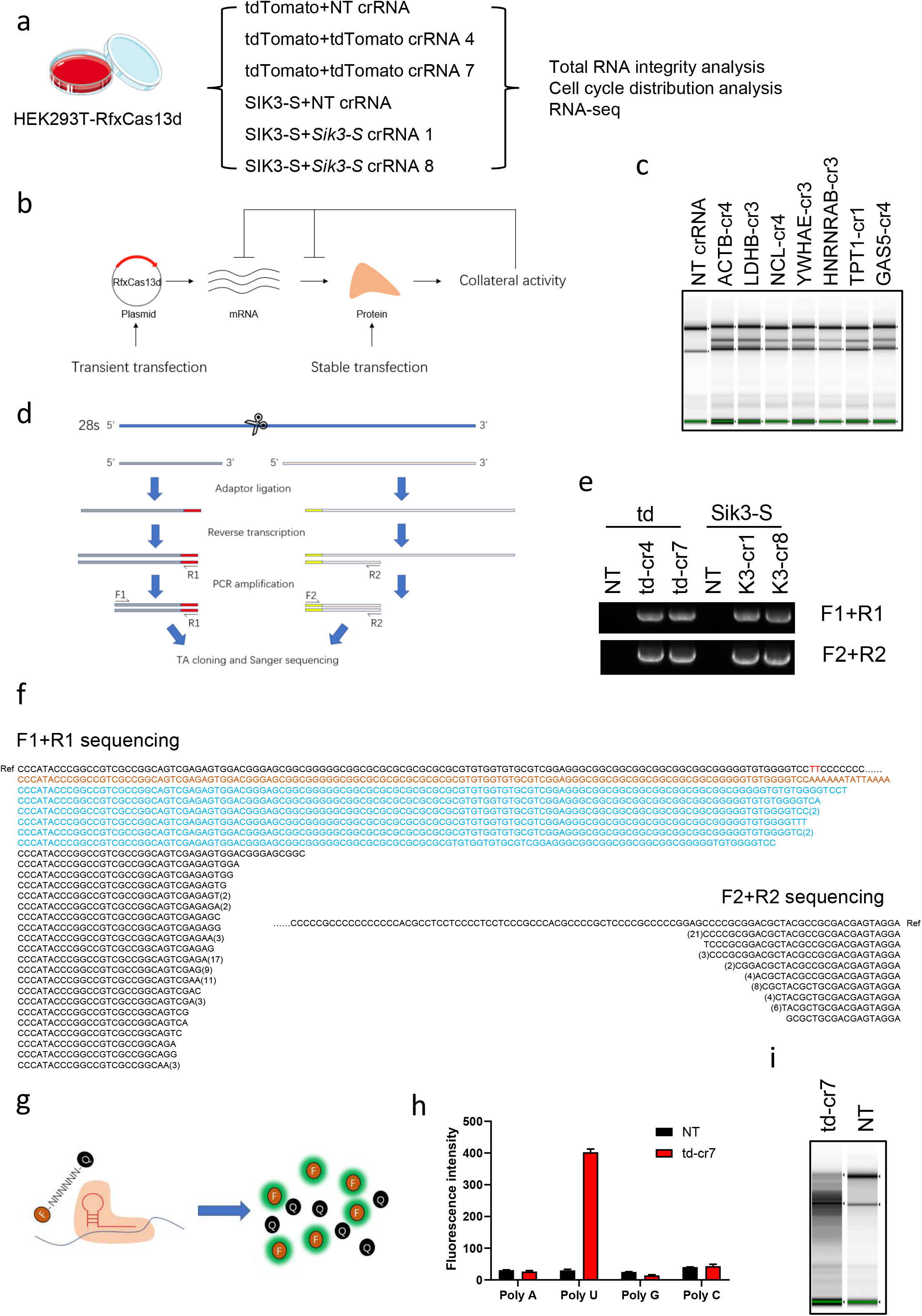
The collateral activity of RfxCas13d cleaves 28s rRNA into two fragments, leading to translation attenuation and activation of ZAKα-JNK/p38-IEG pathway. Linked to Figure 5. a. Workflow of experiments did in HEK293T-RfxCas13d cells. b. Schematic illustration that the collateral activity of RfxCas13d cleaves its own mRNA and 28s RNA, thereby inhibiting its own protein expression. c. Total RNA of each sample was quantified by Agilent 2200 Bioanalyzer. cr: crRNA. d. Schematic illustration of oligonucleotide extension essay. Red represents oligonucleotide adaptor 1; Yellow represents oligonucleotide adaptor 2. e. Gel picture of PCR products from d. f. Sanger sequencing results. Ref: reference sequence. The numbers in brackets represent the number of identical sequencing results. g. Schematic illustration of reconstitution of the collateral activity of RfxCas13d *in vitro*. h. Statistical diagram of fluorescence intensity in g. i. Total RNA integrity analysis quantified by Agilent 2200 Bioanalyzer.

**Figure S6.**
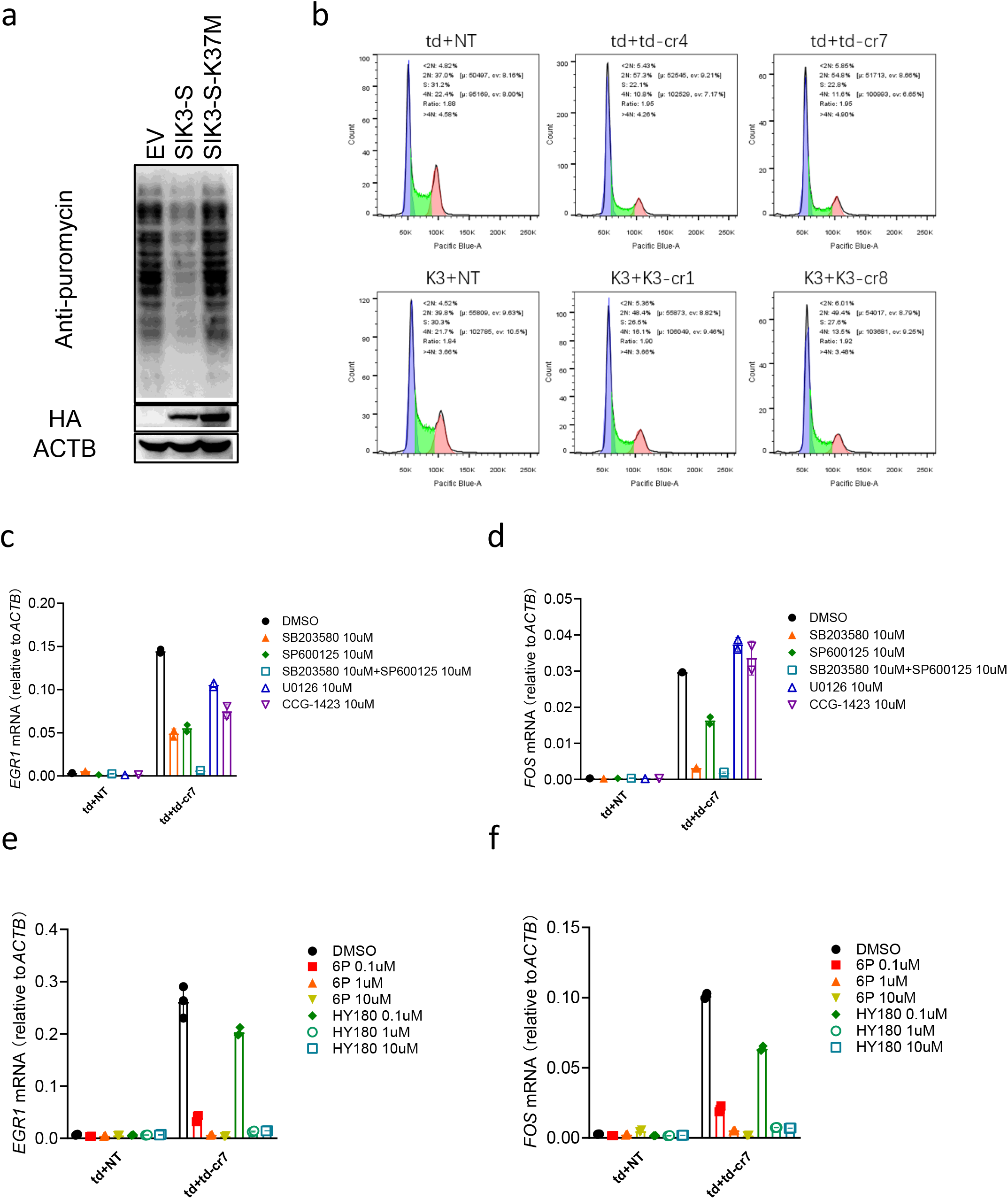
The collateral activity of RfxCas13d cleaves 28s rRNA into two fragments, leading to translation attenuation and activation of ZAKα-JNK/p38-IEG pathway. Linked to Figure 5. a. SUnSET essay to measure the protein translation rate of HEK293T-RfxCas13d cells 24 h after transfection of plasmids encoding SIK3-S or SIK3-S-K37M. b. Cell cycle analysis using FCAS. c-d. RT-qPCR to measure the RNA level of EGR1 and FOS in HEK293T-RfxCas13d cells 24 h after transfection of plasmids encoding tdTomato and crRNAs. Inhibitors and transfection mix were added together. SB203580: p38 inhibitor; SP600125: JNK inhibitor; U0126: MEK1/2 inhibitor; CCG-1423: RhoA/C inhibitor. e-f. RT-qPCR to measure the RNA level of EGR1 and FOS in HEK293T-RfxCas13d cells 24 h after transfection of plasmids encoding tdTomato and crRNAs into. ZAK inhibitors and transfection mix were added together. 6p and HY180: ZAK inhibitors.

## References

1. Makarova KS, Wolf YI, Iranzo J, Shmakov SA, Alkhnbashi OS, Brouns SJJ, et al. Evolutionary classification of CRISPR-Cas systems: a burst of class 2 and derived variants. Nat Rev Microbiol 2020; 18(2):67–83.

2. Shmakov S, Abudayyeh OO, Makarova KS, Wolf YI, Gootenberg JS, Semenova E, et al. Discovery and Functional Characterization of Diverse Class 2 CRISPR-Cas Systems. Mol Cell 2015; 60(3):385–397.

3. Abudayyeh OO, Gootenberg JS, Essletzbichler P, Han S, Joung J, Belanto JJ, et al. RNA targeting with CRISPR-Cas13. Nature 2017; 550(7675):280–284.

4. Meeske AJ, Nakandakari-Higa S, Marraffini LA. Cas13-induced cellular dormancy prevents the rise of CRISPR-resistant bacteriophage. Nature 2019; 570(7760):241–245.

5. Gootenberg JS, Abudayyeh OO, Lee JW, Essletzbichler P, Dy AJ, Joung J, et al. Nucleic acid detection with CRISPR-Cas13a/C2c2. Science 2017; 356(6336):438–442.

6. Cox DBT, Gootenberg JS, Abudayyeh OO, Franklin B, Kellner MJ, Joung J, et al. RNA editing with CRISPR-Cas13. Science 2017; 358(6366):1019–1027.

7. Shmakov S, Smargon A, Scott D, Cox D, Pyzocha N, Yan W, et al. Diversity and evolution of class 2 CRISPR-Cas systems. Nat Rev Microbiol 2017; 15(3):169–182.

8. Konermann S, Lotfy P, Brideau NJ, Oki J, Shokhirev MN, Hsu PD. Transcriptome Engineering with RNA-Targeting Type VI-D CRISPR Effectors. Cell 2018; 173(3):665–676 e614.

9. Yan WX, Chong S, Zhang H, Makarova KS, Koonin EV, Cheng DR, et al. Cas13d Is a Compact RNA-Targeting Type VI CRISPR Effector Positively Modulated by a WYL-Domain-Containing Accessory Protein. Mol Cell 2018; 70(2):327–339 e325.

10. Xu C, Zhou Y, Xiao Q, He B, Geng G, Wang Z, et al. Programmable RNA editing with compact CRISPR-Cas13 systems from uncultivated microbes. Nat Methods 2021; 18(5):499–506.

11. Zhou H, Su J, Hu X, Zhou C, Li H, Chen Z, et al. Glia-to-Neuron Conversion by CRISPR-CasRx Alleviates Symptoms of Neurological Disease in Mice. Cell 2020; 181(3):590–603 e516.

12. He B, Peng W, Huang J, Zhang H, Zhou Y, Yang X, et al. Modulation of metabolic functions through Cas13d-mediated gene knockdown in liver. Protein Cell 2020; 11(7):518–524.

13. Zhou C, Hu X, Tang C, Liu W, Wang S, Zhou Y, et al. CasRx-mediated RNA targeting prevents choroidal neovascularization in a mouse model of age-related macular degeneration. Natl Sci Rev 2020; 7(5):835–837.

14. Wang Q, Liu X, Zhou J, Yang C, Wang G, Tan Y, et al. The CRISPR-Cas13a Gene-Editing System Induces Collateral Cleavage of RNA in Glioma Cells. Adv Sci (Weinh) 2019; 6(20):1901299.

15. Zhang Z, Wang Q, Liu Q, Zheng Y, Zheng C, Yi K, et al. Dual-Locking Nanoparticles Disrupt the PD-1/PD-L1 Pathway for Efficient Cancer Immunotherapy. Adv Mater 2019; 31(51):e1905751.

16. Wang L, Zhou J, Wang Q, Wang Y, Kang C. Rapid design and development of CRISPR-Cas13a targeting SARS-CoV-2 spike protein. Theranostics 2021; 11(2):649–664.

17. Ozcan A, Krajeski R, Ioannidi E, Lee B, Gardner A, Makarova KS, et al. Programmable RNA targeting with the single-protein CRISPR effector Cas7-11. Nature 2021; 597(7878):720–725.

18. Funato H, Miyoshi C, Fujiyama T, Kanda T, Sato M, Wang Z, et al. Forward-genetics analysis of sleep in randomly mutagenized mice. Nature 2016; 539(7629):378–383.

19. Wang G, Li Q, Xu J, Zhao S, Zhou R, Chen Z, et al. 2021.

20. Chan KY, Jang MJ, Yoo BB, Greenbaum A, Ravi N, Wu WL, et al. Engineered AAVs for efficient noninvasive gene delivery to the central and peripheral nervous systems. Nat Neurosci 2017; 20(8):1172–1179.

21. Sasagawa S, Takemori H, Uebi T, Ikegami D, Hiramatsu K, Ikegawa S, et al. SIK3 is essential for chondrocyte hypertrophy during skeletal development in mice. Development 2012; 139(6):1153–1163.

22. Hayasaka N, Hirano A, Miyoshi Y, Tokuda IT, Yoshitane H, Matsuda J, et al. Salt-inducible kinase 3 regulates the mammalian circadian clock by destabilizing PER2 protein. Elife 2017; 6.

23. Wang HY, Hsieh PF, Huang DF, Chin PS, Chou CH, Tung CC, et al. RBFOX3/NeuN is Required for Hippocampal Circuit Balance and Function. Sci Rep 2015; 5:17383.

24. Teng J, Takei Y, Harada A, Nakata T, Chen J, Hirokawa N. Synergistic effects of MAP2 and MAP1B knockout in neuronal migration, dendritic outgrowth, and microtubule, organization. J Cell Biol 2001; 155(1):65–76.

25. Gumucio A, Lannfelt L, Nilsson LN. Lack of exon 10 in the murine tau gene results in mild sensorimotor defects with aging. BMC Neurosci 2013; 14:148.

26. Vicente MM, Chaves-Ferreira M, Jorge JMP, Proenca JT, Barreto VM. The Off-Targets of Clustered Regularly Interspaced Short Palindromic Repeats Gene Editing. Front Cell Dev Biol 2021; 9:718466.

27. Madisen L, Zwingman TA, Sunkin SM, Oh SW, Zariwala HA, Gu H, et al. A robust and high-throughput Cre reporting and characterization system for the whole mouse brain. Nat Neurosci 2010; 13(1):133–140.

28. Vialetto E, Yu Y, Collins SP, Wandera KG, Barquist L, Beisel CL. 2021.

29. East-Seletsky A, O’Connell MR, Knight SC, Burstein D, Cate JH, Tjian R, et al. Two distinct RNase activities of CRISPR-C2c2 enable guide-RNA processing and RNA detection. Nature 2016; 538(7624):270–273.

30. Brogan DJ, Chaverra-Rodriguez D, Lin CP, Smidler AL, Yang T, Alcantara LM, et al. A Sensitive, Rapid, and Portable, CasRx-based Diagnostic Assay for SARS-CoV-2. medRxiv 2020.

31. Bahrami S, Drablos F. Gene regulation in the immediate-early response process. Adv Biol Regul 2016; 62:37–49.

32. Vind AC, Genzor AV, Bekker-Jensen S. Ribosomal stress-surveillance: three pathways is a magic number. Nucleic Acids Res 2020; 48(19):10648–10661.

33. Yang J, Shibu MA, Kong L, Luo J, BadrealamKhan F, Huang Y, et al. Design, Synthesis, and Structure-Activity Relationships of 1,2,3-Triazole Benzenesulfonamides as New Selective Leucine-Zipper and Sterile-alpha Motif Kinase (ZAK) Inhibitors. J Med Chem 2020; 63(5):2114–2130.

